# Spatial patterns of APOBEC mutagenesis in the tumour microenvironment of Asian breast cancer

**DOI:** 10.64898/2026.01.01.697328

**Authors:** Zi Ching Tan, Yang Wu, Zhen Wei Neo, Mai Chan Lau, Soo-Hwang Teo, Joe Poh Sheng Yeong, Siow-Wee Chang, Jia Wern Pan

## Abstract

A germline deletion of the coding region at the *APOBEC3B* (*A3B*) gene that is common in Asian and Oceanic populations has been found to be correlated with increased APOBEC mutagenesis, immune activation, and tumour heterogeneity in breast cancer. The mechanism by which *A3B* deletion and APOBEC mutagenesis induces immune activation in Asian breast cancers is still unclear, but has been thought to involve increased neoantigen burden. In this study, we investigated the mechanisms by which *A3B* deletion and APOBEC mutagenesis may lead to immune activation using Stereo-seq spatial transcriptomics of Malaysian triple-negative breast cancer (TNBC) tumours with and without the *A3B* deletion to examine the spatial expression patterns of *APOBEC3A/B* as well the spatial patterns of APOBEC-associated mutations and neoantigens. Unexpectedly, we found only limited associations between *A3B* deletion, APOBEC-associated neoantigens, and immune activation. Instead, we found strong spatial associations between mitochondrial APOBEC-associations mutations, the cGAS-STING pathway, and the NF-κB pathway. Our study thus supports a new model whereby APOBEC-mediated activation of the immune system does not occur via neoantigen presentation, but instead occurs via mitochondrial hypermutation that drives cGAS-STING activation of the NF-κB pathway. Our study may provide new insights into the spatial landscape of Asian breast tumours and the effect of germline *A3B* deletion and APOBEC mutagenesis on the tumour immune microenvironment.

## Introduction

Apolipoprotein B mRNA editing enzyme catalytic subunit (APOBEC) proteins have been linked to human cancers since its discovery in the late 1990s. For instance, APOBEC cytidine deaminase activity is found in over 75% of breast tumours [1]. *APOBEC1* (A1) and several APOBEC3 (A3) members, who specifically deaminate the cytosine in the TpC motif, can result in mutations in targeted cytosines. Signatures of APOBEC mutagenesis have been detected in over 50% of cancers across multiple cancer types. These signatures can be classified as single-base substitutions (SBS), *kataegis* (local strand-coordinated hypermutation), and *omikli* (diffuse hypermutation). Besides this, APOBEC mutagenesis is associated with subclonal mutations in driver genes and are often observed in metastatic cancers [2, 3], and plays a role in driving treatment resistance in breast cancer [4].

APOBEC3A (A3A), a member of the APOBEC family of proteins, has been implicated as one of the major players involved in APOBEC-associated carcinogenesis. It can induce the formation of tumour-associated mutations by causing DNA strand breaks in somatic cells, triggering the DNA damage response (DDR) and activating DNA replication checkpoints, leading to cell cycle arrest [5]. A3A and its neighbour APOBEC3B (A3B), another member of the APOBEC family, are regarded as the main contributors towards APOBEC mutagenesis. Previous studies suggest that A3A and A3B are the only endogenous enzymes that regularly induce these mutational signatures in human cancer [2]. However, despite generally lower tissue expression than A3B, A3A may have greater enzymatic activity, resulting in more mutations, and these A3A-associated somatic mutations may be more abundant within cancer genomes [6]. APOBEC3-induced mutations are often highly recurrent driver mutation that affect oncogenes and tumour suppressor genes. They can also affect diseases through immune modulation in the tumour microenvironment, by either promoting immune-activated or immunosuppressed cancer phenotypes [6].

The chromosome 22 locus containing *A3A* and *A3B* also contains a common germline polymorphism. This polymorphism is a 29.5 kb germline deletion of the 3’-end of the *A3A* gene and most of the *A3B* gene, producing a hybrid sequence of *A3A* fused with the tail end (3’–untranslated region) of *A3B*. This polymorphism is common in East Asians at a minor allelic frequency of 36.9%, and is also prevalent in Amerindians (57.7%), and Oceanic populations (92.9%). However, the deletion is rarely found in Africans and Europeans (0.9% and 6% respectively) [7]. Translation of the hybrid A3A-B allele results in an identical protein as A3A, but the A3A-B isoform was found to be more stable than the wild-type protein, resulting in higher intracellular levels of A3A [8]. Studies have found that carriers of the germline *A3B* deletion in The Cancer Genome Atlas (TCGA) breast cancer cohort are twice as likely to develop cancers with large numbers of mutations that correspond to the APOBEC mutational signature [9]. Using sequencing data from a Southeast Asian breast cancer cohort [10], we previously found that the polymorphism results in the expression of the A3A-B isoform leading to increased APOBEC-associated somatic hypermutation. APOBEC somatic hypermutation is in turn associated with higher neoantigen burden levels, immune cell presence, tumour heterogeneity, and overall better survival [11].

A3A and A3B are responsible for APOBEC mutations in the tumour epithelium, but studies have found that they are also expressed in immune cells [12]. Furthermore, APOBEC mutagenesis has been linked to immune activation across different cancers [11, 13–17]. We thus sought to study the spatial expression and activity of A3A and A3B in both immune cells and the tumour epithelium in the tumour microenvironment, in order to investigate the potential mechanisms by which APOBEC mutagenesis can activate the immune system, and how tumour cells may potentially evade the immune system in response.

In this study, we performed Stereo-seq spatial transcriptomic analysis on Malaysian triple-negative breast cancer (TNBC) tumours and compared the spatial patterns of APOBEC gene expression, APOBEC-associated mutations, and immune cell types in samples with and without *APOBEC3B* germline deletion. As part of this analysis, we also developed a pipeline to determine the spatial location of single nucleotide variants from spatial transcriptomics data. Our data did not support the hypothesis that APOBEC-mutagenesis leads to immune activation via neoantigen presentation, but instead supported a model whereby APOBEC-associated mutagenesis of mitochondrial DNA led to immune activation via activation of the cGAS-STING and NF-κB pathways. This study may provide insights into the spatial landscape of Asian breast tumours and the effect of germline *A3B* deletion on the tumour microenvironment.

## Results

### Malaysian TNBCs are spatially heterogeneous

Eight Malaysian triple-negative breast cancer (TNBC) tumour samples were selected to undergo Stereo-seq spatial transcriptomics. Among the eight samples, three samples were homozygous for germline *A3B* deletion (A3Bdel), two samples were heterozygous, and three samples were wild-type for the polymorphism (**Table 1**). The samples were given sample names that represent their A3Bdel status: HOM for homozygous, HET for heterozygous, WT for wild-type. The samples, sequenced in pairs, were separated and processed individually. The resulting spatial transcriptomics data (**Figure 1A**, “total counts”) corresponds well to the H&E (**Figure 1A**, “H&E”), with the tumour regions generally having higher gene counts than stromal regions. Cell type deconvolution was performed according to spatial square “bins” (a square 50 µm × 50 µm area) and the resulting cell type scores for each bin were grouped and visualized according to three main categories: immune cells, cancer-associated fibroblasts, and cancer epithelium (**Figure 1A**, “cell types”), revealing significant heterogeneity in the prevalence of different cell types across our samples. Decomposition of these major cell types into more specific minor cell types revealed additional heterogeneity in the tumour microenvironments of our samples (**Supp Figure 1A**).

**Figure 1.**
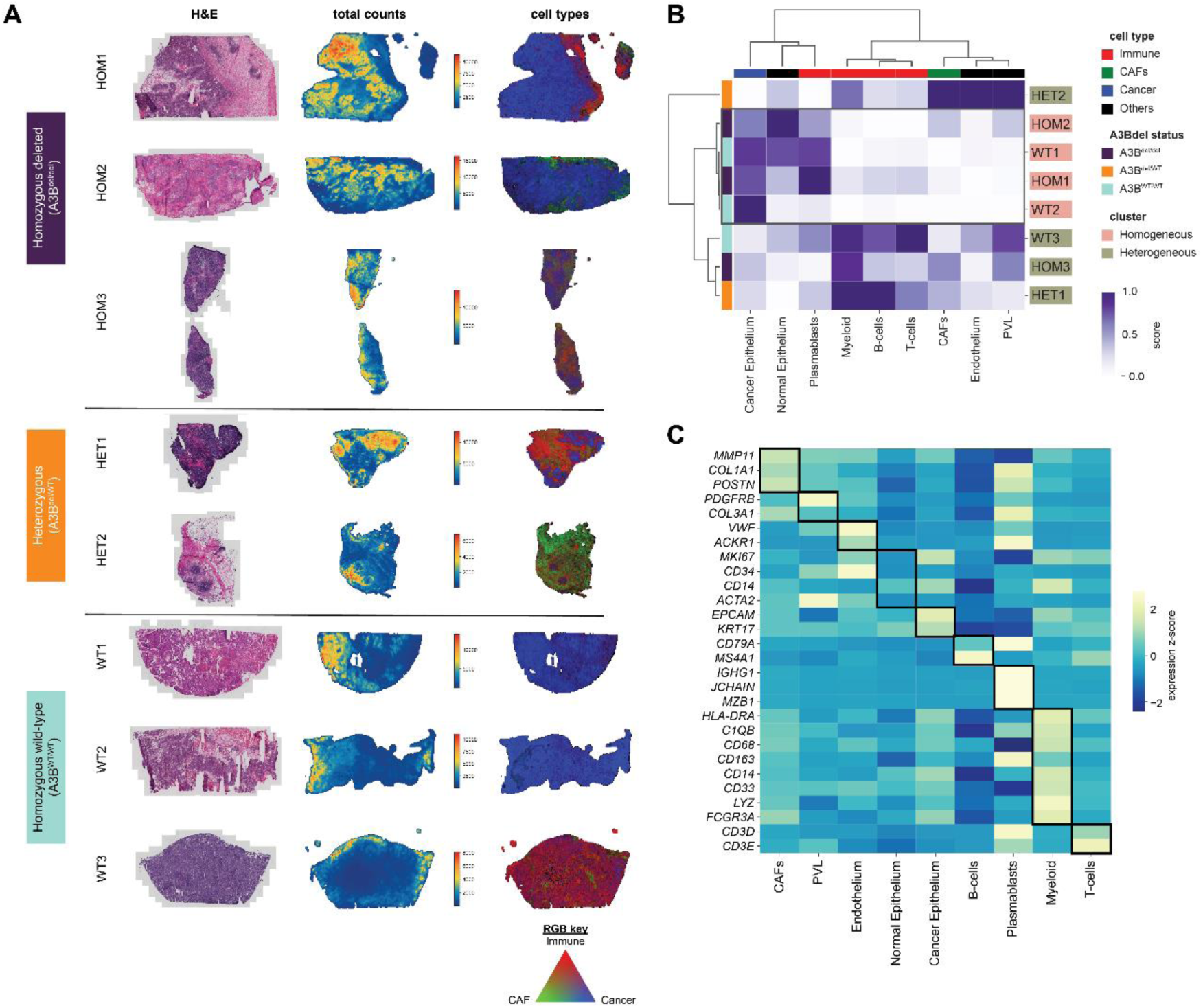
Overview of MyBrCa TNBC samples, clustering and gene expression analyses for major cell types. A: Overview of 8 MyBrCa TNBC samples, first of their H&E staining, followed by number of total transcript counts across each bin, and lastly the composition of immune (red), cancer-associated fibroblast (green), and cancer (blue) cells resulting from cell type deconvolution. B: Unsupervised hierarchical clustering of all samples using major cell type scores resulting from cell type deconvolution. The scores for each sample were first adjusted to a total of 1 prior to clustering. C: Heatmap showing the gene expression score for lineage markers across major cell types. Each bin was assigned a cell type based on the highest score prior to analysis.

**Table 1.**

| Sample name | Sample ID | PAM50 subtype | Clinical subtype | Histology | Age at diagnosis | Ethnicity | Stage | Grade | APOBEC3B deletion status |
| --- | --- | --- | --- | --- | --- | --- | --- | --- | --- |
| **HOM1** | SD0507 | Basal | TNBC | IDC | 56 | Chinese | 2 | 3 | Homozygous deleted (A3B<sup>del/del</sup>) |
| **HOM2** | SD0560 | Basal | TNBC | IDC | 52 | Chinese | 3 | 3 |  |
| **HOM3** | SD1043 | Basal | TNBC | IDC | 49 | Malay | 2 | 3 |  |
| **HET1** | SD1182 | Basal | TNBC | IDC | 31 | Chinese | 2 | 3 | Heterozygous (A3B<sup>del/WT</sup>) |
| **HET2** | SD1225 | Basal | TNBC | IDC | 57 | Chinese | 2 | 3 |  |
| **WT1** | SD0683 | Basal | TNBC | IDC | 45 | Other | 2 | 3 | Homozygous wild-type (A3B<sup>WT/WT</sup>) |
| **WT2** | SD0693 | Basal | TNBC | IDC | 39 | Chinese | 2 | 2 |  |
| **WT3** | SD0781 | Basal | TNBC | IDC | 39 | Chinese | 2 | 3 |  |

Hierarchical clustering of the major cell types across our samples resulted in two distinct clusters, which we labelled as “homogeneous” and “heterogeneous” (**Figure 1B**). The “homogeneous” cluster, composed of two A3B^del/del^ and two A3B^WT/WT^ samples, were composed largely of cancer epithelium, with relatively fewer stromal and immune cells in the tumour microenvironment. Compared to the “heterogeneous” samples, the “homogeneous” samples also contain elevated scores for normal epithelium cells and plasmablasts. The “heterogeneous” cluster, composed of one A3B^del/del^, two A3B^del/WT^ and one A3B^WT/WT^ samples, had more diverse tumour microenvironments with enrichment of various immune cells as well as relatively lower scores for cancer and normal epithelial cells. Hierarchical clustering using minor cell types (**Supp Figure 1B**) refined the previously defined “heterogeneous” cluster, separating HET2 from the other members of the cluster, but the clustering results otherwise remained the same. Overall, our samples were highly heterogeneous with distinct cellular compositions and tumour microenvironments.

To validate the cell type assignments, we assigned each bin its top-scoring cell type, and compared the expression of lineage markers corresponding to each major cell type (**Figure 1C**). High expression of cell type specific lineage/classical markers were found in the corresponding cell types, validating the cell type deconvolution calls. For further validation, we used a pseudobulk method to generate cell type scores for each sample by averaging the cell type scores across all bins in a sample. The pseudobulk scores were then compared with ssGSEA scores obtained from matched bulk RNA-seq data (**Supp Figure 1C**). Positive Pearson correlation between the pseudobulk and bulk RNA-seq scores indicated that our cell type deconvolution results were reliable.

### Spatial patterns of APOBEC gene expression in Malaysian TNBCs

We then investigated the expression of APOBEC genes, particularly *APOBEC3A* and *APOBEC3B*, the two genes that are the most affected by the *A3B* deletion polymorphism (**Figure 2A**). The expression of these genes was sparse and unevenly distributed across the tissue, especially in comparison with the expression patterns of housekeeping genes (**Figure 2A**, **Supp Figure 2**). We then compared the expression of APOBEC and housekeeping genes across all spatial bins based on their A3Bdel status (presence or absence of the germline *A3B* deletion in the sample that the bin is from), and found that expression of both *A3A* and *A3B* were significantly lower in A3Bdel carriers compared to non-carriers (**Figure 2B**). Due to concerns that this finding may be attributed to zero-inflation in the expression scores, we also made the same comparison using only bins with non-zero expression of the *A3A* and *A3B* genes, and found that in these bins, *A3A* expression was significantly higher in A3B^del/del^, while most A3B^del/del^ bins had no *A3B* expression except for a few outliers (**Figure 2C**). Additionally, bins expressing *A3A* and *A3B*, although rarer, had higher expression scores in those bins compared to housekeeping genes (**Figure 2C**). These data suggests that *A3A* and *A3B* are only expressed in a subset of cells, but when they are expressed, they are expressed at a high level. We also compared the expression of APOBEC genes to our cell type scores for each bin, and found that *A3A* and *A3B* had the highest expression in the cancer epithelium relative to other cells type (**Figure 2D**).

**Figure 2.**
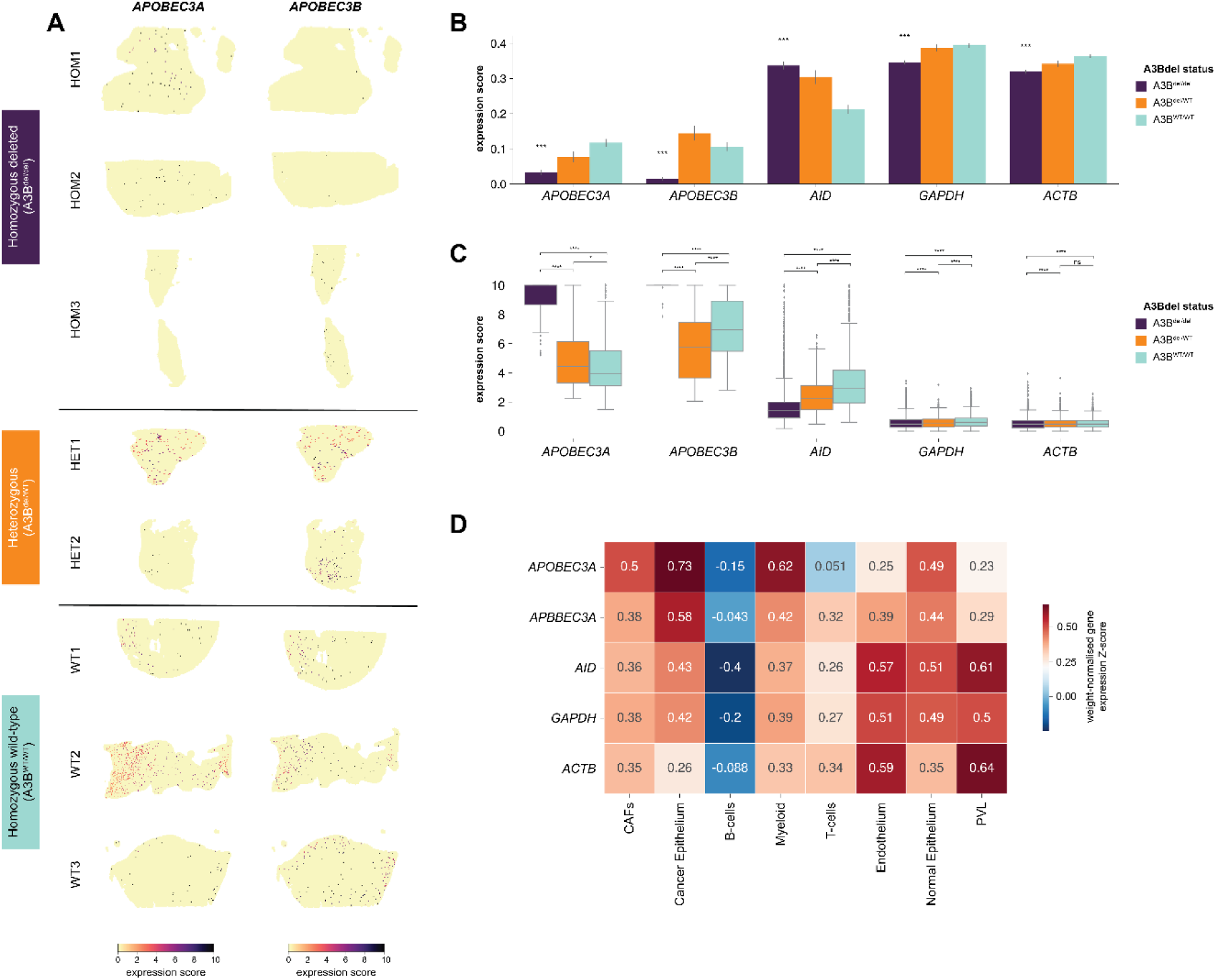
Expression of APOBEC and housekeeping genes. A: The spatial expression of *APOBEC3A* and *APOBEC3B*. The scores were normalised and scaled. B: Barplots showing mean gene expression scores of *APOBEC3A*, *APOBEC3B*, with housekeeping genes as controls across different A3Bdel statuses. Stars indicate significance (Kruskal–Wallis H test < 0.001) when comparing A3Bdel carriers to non-carriers. C: Boxplots showing the distribution of non-zero gene expression scores of *APOBEC3A*, *APOBEC3B*, with housekeeping genes as controls across different A3Bdel statuses. P-values indicated are for Kruskal-Wallis H test. D: Heatmap showing cell type weight-normalised gene expression Z-scores of *APOBEC3A*, *APOBEC3B*, with housekeeping genes as controls across major cell types. Plasmablasts were excluded from this analysis as there were very few plasmablast bins.

### Spatial patterns of APOBEC-associated mutations in Malaysian TNBCs

Next, we extracted somatic single nucleotide variants (SNVs) from the Stereo-seq data and mapped them to the spatial coordinates of their corresponding transcripts. In order to confirm that these variants are biologically meaningful, we compared the Stereo-seq variants to somatic SNV calls from bulk tumour whole-exome (WES) and bulk tumour RNA-sequencing from the same tumour (but, importantly, from different tumour specimens). After filtering for high quality variants (supported by at least 10 reads in our Stereo-seq data), of the somatic SNVs that were found in both bulk tumour WES and bulk tumour RNA-seq, 88.4% of those SNVs were also found in our Stereo-seq data across all samples (**Supp Table 1-2**). The high concordance between bulk WES, bulk RNA-seq, and Stereo-seq suggests that Stereo-seq captures variants that are biologically relevant, although there was some variability between samples that was likely due to differences in tumour heterogeneity (**Supp Table 2**). We also compared a random subset of germline variants across all sequencing modalities and found a concordance rate of 96% for germline variants between Stereo-seq and bulk sequencing (**Supp Table 3**).

Closer inspection of the somatic SNVs from Stereo-seq revealed that many of the most prevalent mutations were present in immune-related genes, which was particularly prominent in two samples (**Supp Figure 3A**). For example, we noted a large number of mutated transcripts in the *IGHG4* gene in one sample (HET1) that originated from a common single point mutation (c.974T>C), and in the same sample we also found that areas with high mutation scores (areas containing high normalized counts of variant-containing transcripts) also had high immune and stromal cell type scores, suggesting that many of these variants are likely associated with immune function. We also performed a spatial analysis of tumour mutation burden (spatial TMB) by quantifying the number of unique variants found in each bin (**Supp Figure 3B**) and found that TMB was highest in spatial regions corresponding to the tumour epithelium. Together, these data demonstrate that the variants captured by Stereo-seq are likely to be biologically relevant, and that it is possible to distinguish between somatic variants in tumour cells versus immune/stromal cells.

We next focused specifically at the spatial distribution of Stereo-seq SNVs associated with APOBEC mutational signatures, which were present in all samples but were unevenly distributed (**Figure 3A**). We defined mutational signatures in this study based on the most common trinucleotide context for each mutational signature as defined in the COSMIC library of mutational signatures. Using this definition, SBS1-associated variants had a broad distribution across samples while SBS10a-associated variants had a relatively sparse distribution similar to APOBEC-associated SNVs (**Supp Figure 4A**). When SBS2- and SBS13-associated mutations, which make up the APOBEC signature, were examined separately, we found that our samples generally had a higher prevalence of SBS2 than SBS13 (**Figure 3B**). We also compared specifically YT<u>C</u>A (A3A-associated) and RT<u>C</u>A (A3B-associated) mutations between A3Bdel carriers versus non-carriers, which showed that carriers had more YT<u>C</u>A and less RT<u>C</u>A mutations, and thus more variants associated with the activity of the A3A enzyme (**Figure 3**C). Next, we quantified the somatic variants assigned to various signatures and compared them by the A3Bdel status of the bins in which they were located. We found that A3B^del/del^ had significantly higher APOBEC mutation scores compared to A3B^del/WT^ and A3B^WT/WT^. This was found to be driven mainly by SBS2, rather than SBS13. However, we also found that the distribution of RNA-editing SBS2-associated variants did not correlate with A3Bdel status (**Figure 3**D). On this point, we also found significant heterogeneity among samples for mutation scores for each mutational signature, so these results should be interpreted cautiously (**Supp Figure 4**B). Nonetheless, SBS2 and SBS13-associated variants were located primarily within the cancer epithelium, as expected (**Figure 3**E, **Supp Figure 5**).

**Figure 3.**
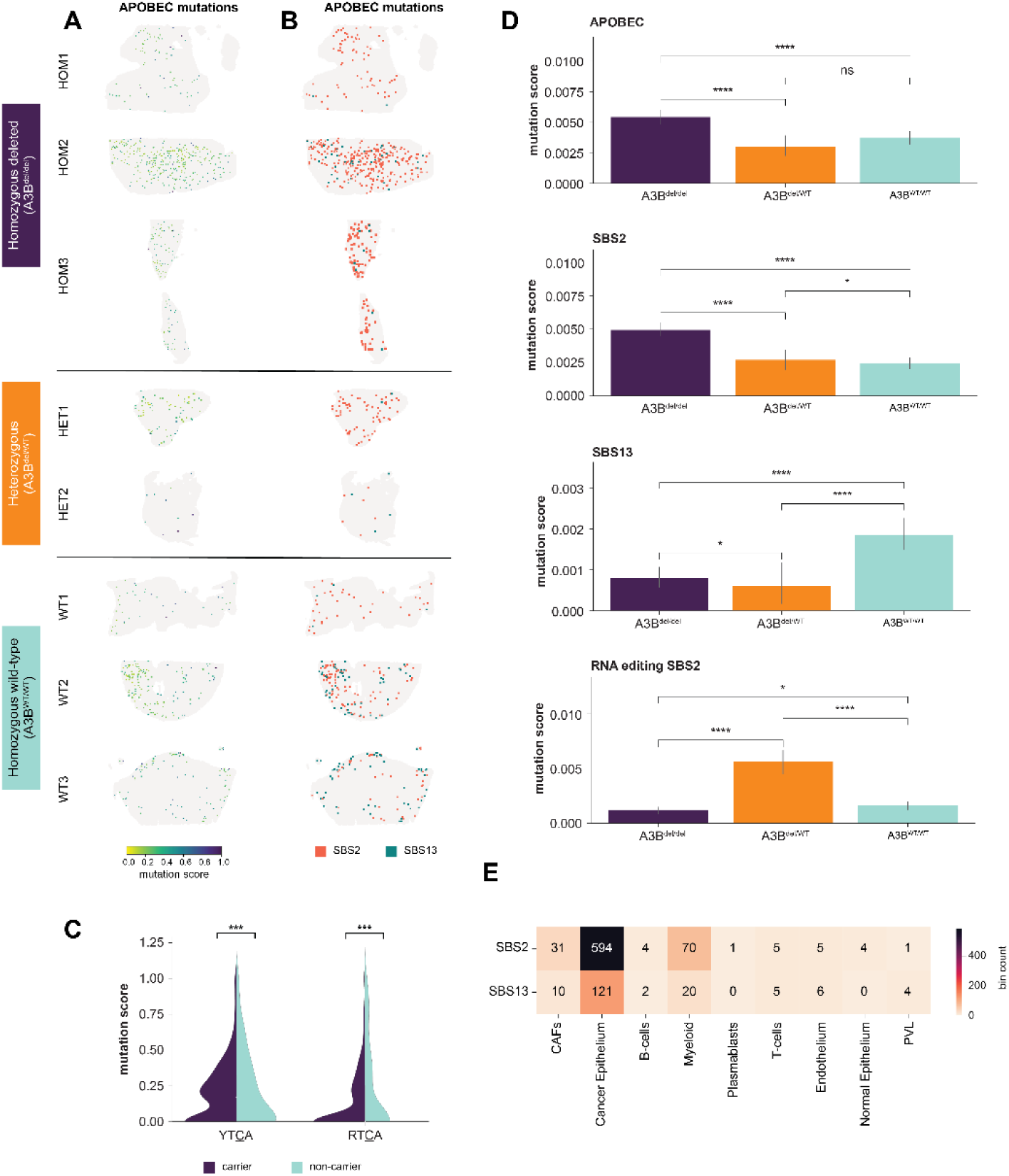
Distribution of somatic APOBEC-associated single nucleotide variants (SNVs). A: Spatial distribution of APOBEC-associated SNV mutation scores across all samples. Mutations scores were mutation counts normalised against total transcript counts in each bin. B: Spatial distribution of SBS2 and SBS13 mutations across all samples. C: Split violinplots showing the distribution of A3A-associated YT<u>C</u>A mutations and A3B-associated RT<u>C</u>A mutations in carriers versus non-carriers. P-values indicated are for Mann-Whitney U test. D: Barplots showing mean mutation scores for somatic APOBEC, SBS2, SBS13 and RNA-editing SBS2 mutations across different A3Bdel statuses. P-values indicated are for Mann-Whitney U test. E: Heatmap for the number of bins containing SBS2 and SBS13 mutations. Each bin was assigned a cell type based on the highest score prior to analysis.

### Spatial patterns of neoantigens in Malaysian TNBC

Following our analysis of Stereo-seq somatic SNVs, we computationally predicted which of these SNVs would result in neoantigens, and mapped these predicted neoantigens to their corresponding spatial bins. We observed predicted neoantigens in all the samples, with significant heterogeneity in their spatial distribution (**Figure 4**A, **Figure 4**B). We found little to no significant difference in neoantigen scores (normalized quantification of predicted neoantigen load) across bins with different A3Bdel status, suggesting that there is no association between germline *A3B* deletion and neoantigen formation (**Figure 4**C). The predicted neoantigens were also predominantly found in the cancer epithelium, as expected (**Figure 4**D, **Supp Figure 6**A). Next, we also looked at the trinucleotide context of the predicted neoantigens in order to determine which mutational signatures were more likely to generate neoantigens. We found that the predicted neoantigens present in the cancer epithelium originated mainly from C>T, C>G and T>C mutations (**Figure 4**E), with C[C>T]G derived predicted neoantigens from the *PSENEN* gene being particularly concentrated/dominant in one sample (**Supp Figure 6**B). Importantly, the variants associated with the APOBEC mutational signatures SBS2 and SBS13 did not seem to generate many predicted neoantigens (Figure 4E). In other cell types aside from cancer epithelial cells, C>T and T>C transitions were the mutations that were most likely to generate predicted neoantigens, with T>C mutations appearing particularly consistent in importance across the cell types (**Supp Figure 7**).

**Figure 4.**
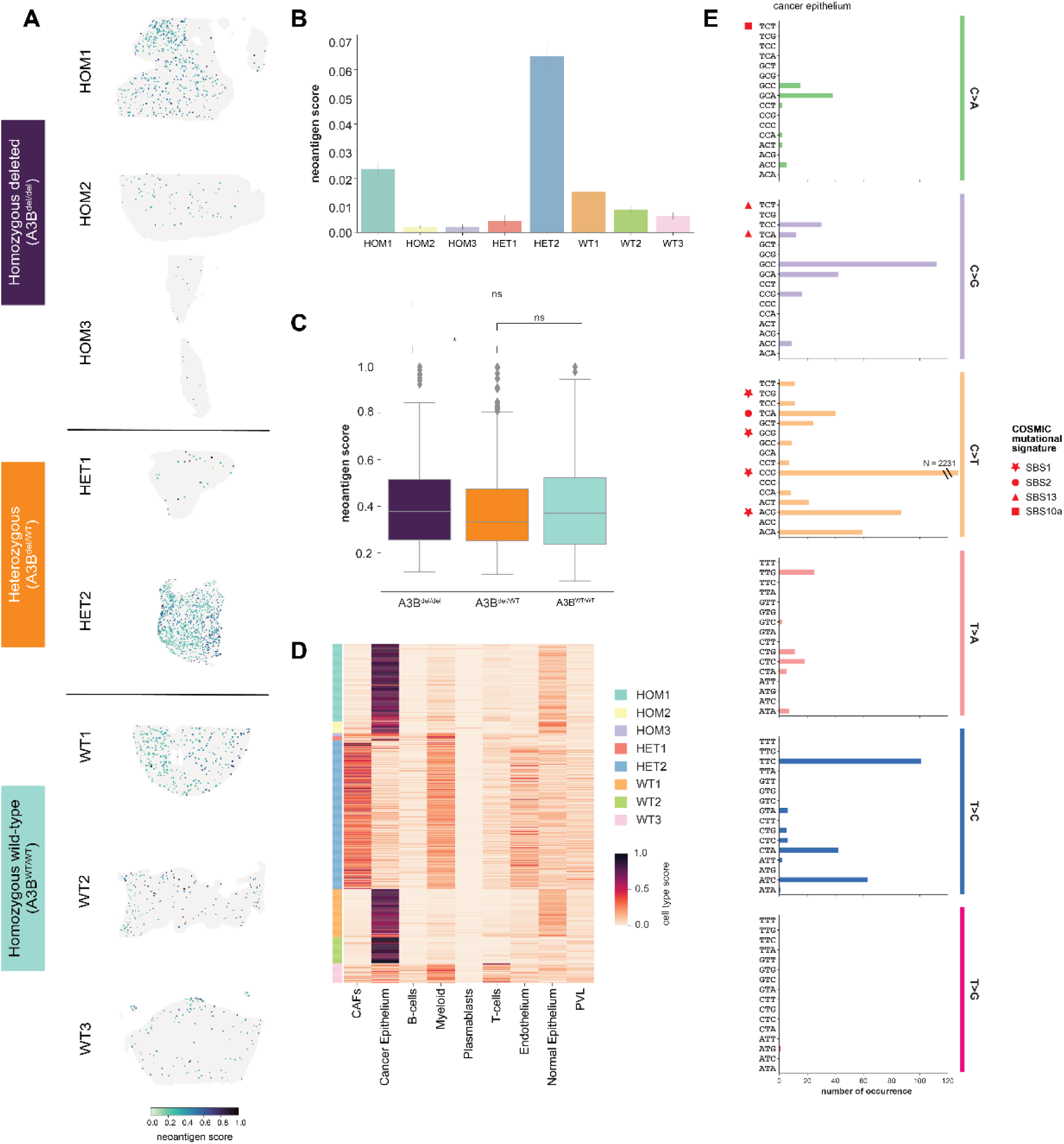
Spatial distribution of candidate neoantigens. A: Spatial distribution of candidate neoantigen scores across all samples. B: Barplots showing the average neoantigen scores across all samples. C: Boxplots showing the distribution of non-zero neoantigen scores across different germline A3Bdel statuses. P-values indicated for Mann-Whitney U test. D: Heatmap showing major cell type scores for bins predicted of having neoantigens across all samples. E: Quantification of SNVs, specifically single base substitutions, predicted to give rise to neoantigens, limited to bins previously determined to be cancer epithelium. The scale has been truncated, and values exceeding the scale are annotated within the figure. Mutational signatures of interest are highlighted.

### Immune activation in TNBC may be countered by tumour resistance

Next, we examined the spatial distribution of immune cells in our samples, dividing the immune cell types into four main categories (plasmablasts, B-cells, T-cells, and myeloid cells) and assigning each of them a colour in the CMYK colour scheme according to their cell type scores (**Figure 5A**). Visualisation of the tumour immune microenvironment in this way revealed the presence of tertiary lymphoid structures (TLS) in three samples (HOM1, HOM3, HET1), with TLS being defined as clusters of B-cells surrounded by T-cells. The TLS were particularly prominent in one sample (HOM1), and a deeper look at the TLS in this sample showed presence of both naïve and memory B-cells along with *CCR7*+ CD4+ T-cells and *CXCL13*+ T follicular helper (Tfh) cells (**Supp Figure 8**). In addition, we also noted that one sample (WT3) was an outlier among our samples, with remarkable levels of immune infiltration (**Figure 5**A, **Figure 1**A). Hierarchical clustering of the immune and CAF cell types across samples recapitulated the “homogeneous” cluster from our previous analysis, but the “heterogeneous” cluster was further split into 2 subclusters, which we designated as “immune-enriched” and “CAF-enriched” subclusters (**Figure 5**B). A closer look at various immune cell subtypes (dendritic cells, T-cells, B-cells) revealed heterogeneity among samples with no clear trend associated with A3Bdel status (**Figure 5**C, **Supp Figure 9**A, **Supp Figure 9**B).

**Figure 5.**
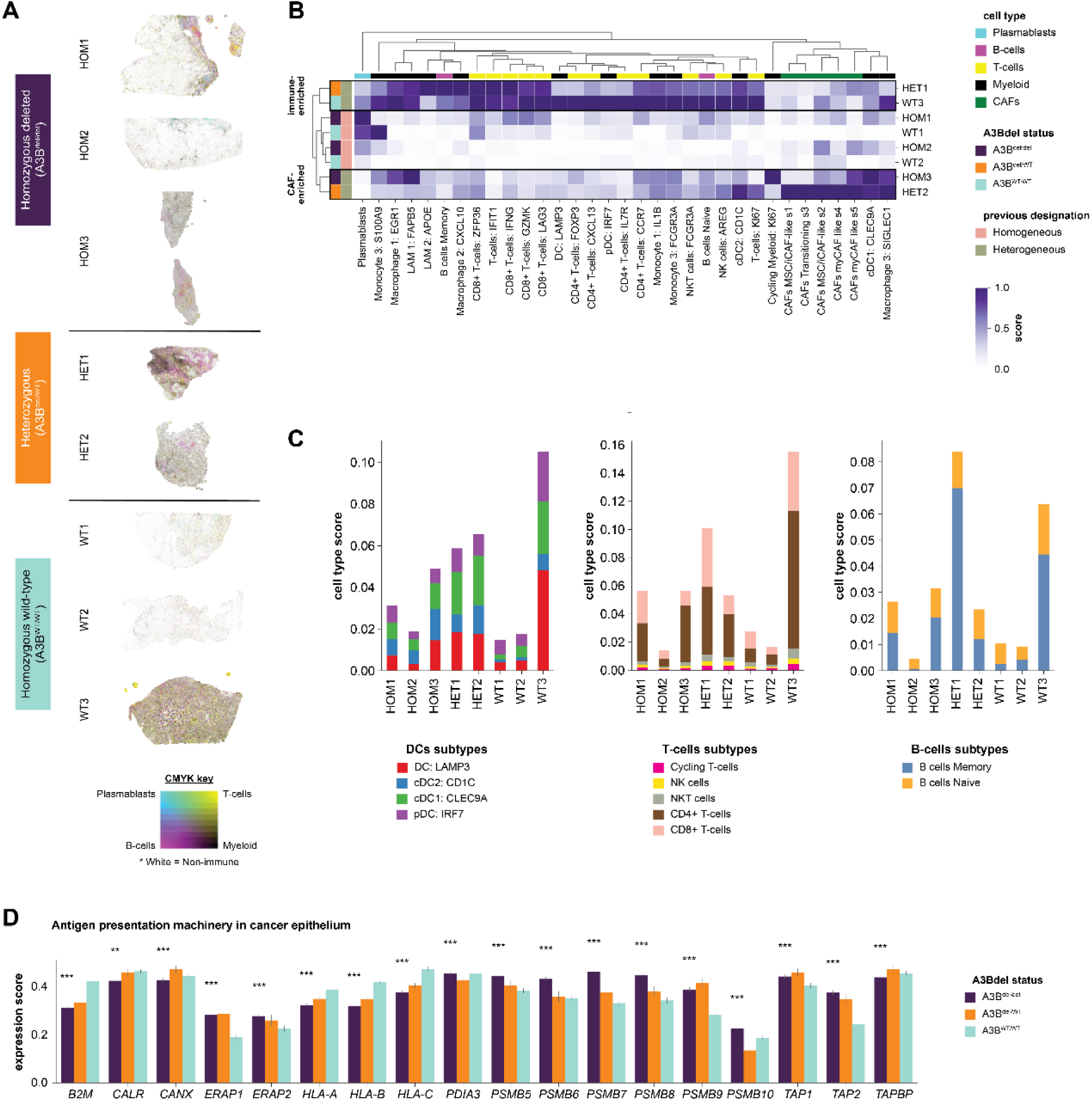
The tumour immune landscape of TNBC. A: Composition of the tumour immune microenvironment of TNBC, broken down into plasmablasts (cyan), B-cells (magenta), T-cells (yellow), and myeloid cells (black). B: Unsupervised hierarchical clustering of all samples using minor cell type scores for immune and CAF cell types. The scores for each sample were first adjusted to a total of 1 prior to clustering. C: Barplots showing the average cell type scores for DCs, T-cells, and B-cells, each broken down into individual subtypes, across samples. D: Average gene expression scores for genes in the antigen presentation machinery across different germline *A3B* deletion statuses.

We then looked at the expression of genes belonging to the antigen presentation machinery pathway, and observed significant differences in expression associated with A3Bdel status, notably in the *HLA* genes, which were downregulated in carriers, and *TAP* genes, which were upregulated in the carriers (**Figure 5**D). However, when the data was filtered only for bins with non-zero expression, the opposite trend was observed for many of these genes, suggesting an inverse relationship between sparsity and expression level, thus demonstrating the caveats of analysing zero-inflated spatial data (**Supp Figure 9**C).

Broadly speaking, we found that A3Bdel carriers generally had more inflamed tumour microenvironments; however, immune infiltration was not a guarantee in these samples. For example, samples HOM1 and HOM2, both of which were homozygous for A3Bdel, contained evidence of immune cell recruitment in the periphery of the tumour bed but had little to no immune infiltration into the tumour bed, representing an “immune-excluded” phenotype (**Figures 1A, 5A**). For these samples, we hypothesized that there were additional factors present that were suppressing the infiltration of these immune cells into the tumour bed in these samples. Upon further investigation, we found that these samples had hotspots of *IDO1* expression, a known immunosuppressive gene [18]. However, the spatial expression of *IDO1* overlapped with *CXCL10*, a pro-inflammatory cytokine (Pearson correlation with *IDO1* expression = 0.26 and 0.23, respectively), suggesting a complex interplay between pro- and anti-inflammatory factors in the microenvironment of these tumours (**Supp Figure 10**A,B,C). Additionally, in one of these samples (HOM2), we further observed a layer of CAFs, determined to be myofibroblastic CAFs (myCAFs), lining the edge of the tumour (**Figures 1A, Supp Figure 10**D), which may have acted as a physical barrier to potentially hinder the movement of immune cells into the tumour [19, 20].

### Association between APOBEC mutagenesis and activation of the cGAS-STING pathway in Malaysian TNBCs

Given that we found little association between APOBEC mutagenesis and predicted neoantigens, we hypothesised that APOBEC-associated mutagenesis (resulting in part from germline *A3B* deletion) may instead be driving immune activation by activating the cGAS-STING pathway through generating oxidative stress on the mitochondria [5, 21–27]. To test this hypothesis, we measured the correlation between different components in this pathway, including C>T mitochondrial DNA (MT) mutations, *APOBEC3A*, *NOX4*, *CGAS*, and *NFKB1*, as well as positive and negative controls, limiting our analysis to the cancer epithelium (**Figure 6**A). We also limited our analyses to bins with non-zero scores for both variables in order to account for zero-inflation in the data. These analyses showed that, within the bins predicted to be cancer epithelium, *A3A* expression was moderately correlated with C>T MT mutations (Pearson correlation = 0.31, **Figure 6**A). The C>T MT mutations were in turn strongly correlated with *NOX4* expression (Pearson correlation = 0.57, **Figure 6**A), indicating a positive relationship between APOBEC-associated MT mutations and ROS production. *NOX4* expression was in turn strongly correlated with *CGAS* expression (Pearson correlation = 0.62, **Figure 6**A), linking ROS to increased presence of cytosolic DNA. Finally, we found that *CGAS* was in turn correlated with *NFKB1* (Pearson correlation = 0.56, **Figure 6**A), completing the known sequence of this immune pathway (**Figure 6**A). As additional validation, we found strong positive correlation between *NFKB1* and *NOX4* (Pearson correlation = 0.68, **Figure 6**A) as well as *CGAS* and C>T MT mutations (Pearson correlation = 0.57, **Figure 6**A). As a negative control, we found much weaker correlations between *GAPDH* and C>T MT mutations as well as between *GAPDH* and *NFKB1* (Pearson correlation = 0.12 and 0.05, respectively; **Figure 6**A). We also examined the gene expression of immune-related genes downstream of the cGAS-STING pathway, and found that genes associated with ROS, Type I interferon (IFN) and its associated chemokines were all upregulated in carriers of germline *A3B* deletion (**Figure 6**B). Taken together, our results are consistent with a model whereby APOBEC mutagenesis of mitochondrial DNA leads to immune activation via cGAS-STING mediated upregulation of the NF-κB pathway (**Figure 6**C).

**Figure 6.**
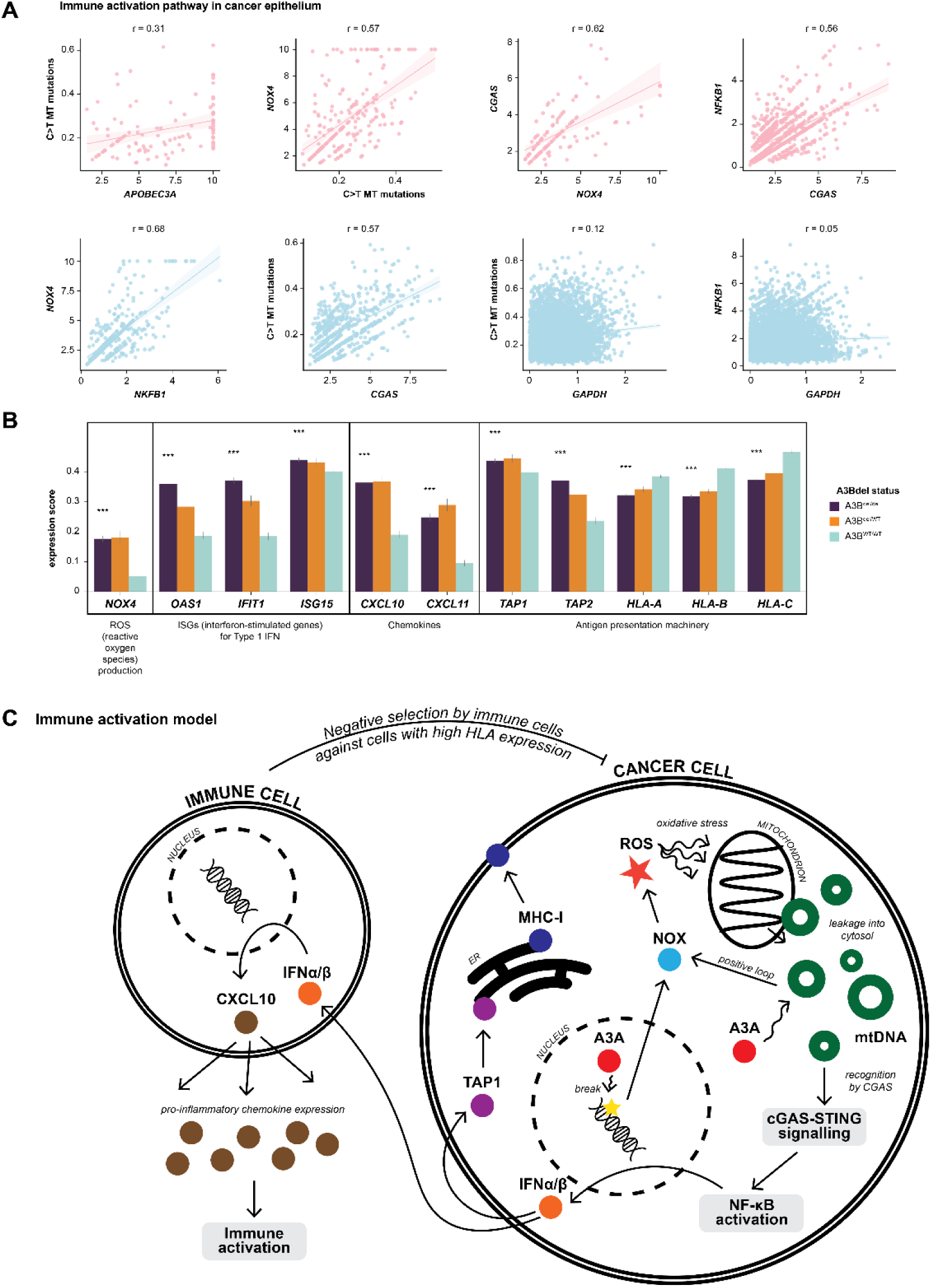
Model of immune activation in through APOBEC-associated mutagenesis. A: Pearson correlation between elements in our hypothesised immune activation pathway: C>T mitochondrial (MT) mutations, *APOBEC3A*, *NOX4*, *CGAS*, and *NFKB1*. Validations are included for additional correlations along the hypothesised pathway, and *GAPDH* as negative control. B: Average gene expression scores for genes involved in reactive oxygen species (ROS) production, Type 1 interferon activation, chemokines, and part of the antigen presentation machinery, compared across different germline *A3B* deletion statuses. C: Illustration of our model of immune activation. The model begins in the nucleus where A3A induces DNA damage, which releases ROS that induces oxidative stress on the mitochondrion, leading to the release of mtDNA into the cytosol, subsequently activating the cGAS-STING and NF-κB pathway, ultimately resulting in immune activation.

## Discussion

In this study, we examined the relationship between germline *A3B* deletion, APOBEC mutagenesis, and the tumour immune microenvironment by analysing Stereo-seq spatial transcriptomics data from eight TNBC samples with varying germline *A3B* deletion status. Cluster analysis of the cell type scores resulted in two clusters of samples with different degrees of homogeneity in cell type compositions. Next, we developed and applied a bioinformatics pipeline to extract somatic SNVs and predicted neoantigens and map them to their corresponding spatial locations. Using these data, we quantified and compared the expression and presence of various genes, mutations and neoantigens across different cell types in the tumour microenvironment. As expected, we found that APOBEC gene expression, APOBEC-associated mutations and predicted neoantigens were localized primarily to the cancer epithelium. We identified significant heterogeneity and complexity in immune cell types across our samples as well as the presence of tertiary lymphoid structures (TLS) in A3Bdel carriers, along with downregulation of *HLA* and upregulation of *TAP* genes in the A3Bdel carriers. Our results also suggested that germline *A3B* deletion and APOBEC mutagenesis may not be associated with neoantigen burden, but instead may activate the immune system through an alternate pathway whereby APOBEC mutagenesis of the mitochondrial genome leads to increased cGAS-STING activation and upregulation of the NF-κB pathway.

Our results demonstrate that the tumour microenvironments of Malaysian TNBCs are extremely heterogeneous, regardless of their germline *A3B* deletion status. Heterogeneity among TNBC tumours has been a known challenge in the quest for developing immunotherapy for TNBC, which does not respond well to conventional therapies [28]. In particular, despite TNBC being typically known for having higher levels of tumour-infiltrating lymphocytes [29–31], we also found significant heterogeneity in immune infiltration and immune cell types among TNBC tumours, which is consistent with previous studies [32, 33].

In carriers of germline *A3B* deletion, we found higher levels of *A3A* expression than *A3B*, consistent with previous studies [6]. In healthy individuals, *A3A* and *A3B* are mainly expressed in the white blood cells of normal tissue (GTEx). In TNBC, we found that both *A3A* and *A3B* are expressed in the cancer epithelium. This is consistent with existing literature, where studies have shown that *A3A* and *A3B* are more highly expressed in the tumour epithelium than the normal epithelium across different cancer types and cell lines [34, 35]. Our results thus confirm that, in TNBC samples, expression of *A3A* and *A3B* occurs primarily in cancer epithelial cells, and less so in immune cells. However, we did observe some expression of both *A3A* and *A3B* in myeloid cells in some samples, specifically in macrophages, which is also consistent with existing reports [5].

With regards to APOBEC-associated mutations, our samples had more A3A-driven YT<u>C</u>A than A3B-driven RT<u>C</u>A mutations, which is consistent with literature reports [11, 36]. APOBEC mutations in our sample, similarly to the expression of *A3A* and *A3B*, were found mainly within the cancer epithelium. This was expected given that the APOBEC mutational signature is a known driver of cancer [37–39]. We also identified more APOBEC mutations in A3B^del/del^ tumours than in A3B^del/WT^ and A3B^WT/WT^ tumours, echoing previous findings [11, 39].

Our pipeline was also able to map predicted neoantigens to their spatial locations. Our analysis showed that the predicted neoantigens were expressed mainly in the cancer epithelium, as expected. We found that there was no difference in neoantigen levels between carriers and non-carriers of the germline *A3B* deletion, and that few APOBEC-associated mutations generated predicted neoantigens. This discovery was surprising to us but aligned with some previous findings [11], suggesting that increased APOBEC mutagenesis does not necessarily lead to higher neoantigen load.

Consistent with previous studies, we found that, despite our small sample size, carriers of the germline *A3B* deletion tended to have more active tumour immune microenvironments. Several of our samples also contained TLS, which has been shown to be a double-edged sword as they can influence the tumour immune microenvironment in either direction [40]. TLS are commonly found in TNBC and can serve as prognostic markers for treatment response [41]; however, it is also important to note that there is also heterogeneity in TLS composition and location, hence their characteristics should be carefully examined in order to maximise their prognostic and therapeutic potential [42, 43].

Some of our samples displayed an “immune excluded” phenotype whereby immune cells where mostly found only on the periphery of the tumour, suggesting that additional mechanisms of immune resistance were in play. In one of these samples, we observed myCAF expression around the tumour; these dense stromal cells may be acting as physical barriers that restrict the movement of immune cells into the tumour [19, 20]. Besides this, *IDO1*, which are sometimes secreted by endothelial cells [19], can aid in suppressing immune activity in breast tumours [18]. This mechanism is supported by some of our data, although additionally complexity may be present.

On the whole, our results are consistent with a model whereby APOBEC mutagenesis of mitochondrial (mt)DNA leads to immune activation via cGAS-STING mediated upregulation of the NF-κB pathway (**Figure 6**C). We have previously shown that germline *A3B* deletion led to increased A3A-mediated mutagenesis [11]. Other studies have found that DNA damage, induced by the activity of A3A deamination can lead to the production of ROS through NADPH oxidase (NOX) enzymes [5, 21, 22]. Excessive ROS can exert oxidative stress on the mitochondria, disrupting the integrity of mtDNA and facilitating the release of oxidised mtDNA into the cytoplasm [23, 24]. Additionally, A3A favours cytosines present in damaged motifs and the activity of A3A increases ssDNA availability [21, 22]. A3A can also deaminate and hypermutate leaked mtDNA within the cytosol [21], which could potentially form a positive loop that ultimately leads to increased levels of cytosolic mtDNA. These cytosolic mtDNA are then recognised by cGAS, a cytosolic DNA sensor, which subsequently activates the cGAS-STING pathway [23–25]. The cGAS-STING pathway first cascades through the cGAS-cGAMP-STING-TBK1-IRF3 axis [26] before activating IKK and subsequently NF-κB, leading to Type I IFN response [24, 26, 27]. Pro-inflammatory chemokines, including CXCL10, are released upon Type I IFN signalling to promote immune activation [26] in the tumour microenvironment. Our analyses of the spatial transcriptomics data recapitulated some of the key associations and correlations suggested by this model. We also included in this model our hypothesis that negative selection by the immune cells may be at play to select against cancer cells with high MHC-I expression, which may explain the lower *HLA* expression that we see in carriers of A3Bdel. Our model thus could explain how germline *A3B* deletion, through increased A3A-driven APOBEC mutagenesis, can lead to immune activation in tumours carrying the polymorphism.

Taken together, our work does not support previous theories of APOBEC-mediated activation of immune pathways, which centred around the generation of neoantigens. Instead, we found more support for APOBEC activation of immune pathways via cGAS-STING, through generation of oxidative stress on mtDNA by APOBEC mutagenesis. However, it is important to note that tumours are highly heterogeneous; each tumour in our dataset had its own unique characteristics and immune microenvironment and was likely shaped by very different factors. Our work ultimately highlights the heterogeneity among TNBCs in general and the need for personalised therapy to target the unique characteristics of each tumour.

## Methods

### Study cohort

Our study cohort consists of breast cancer samples from the Malaysian Breast Cancer Genetic Study (MyBrCa), that were recruited from Subang Jaya Medical Centre and University Malaya Medical Centre in Kuala Lumpur, Malaysia between 2012 and 2018 [10, 44]. Representative fresh breast tumour tissues that were obtained during surgical resection of the tumour were flash frozen and then stored in liquid nitrogen.

Triple-negative breast cancer (TNBC) samples were selected from the MyBrCa cohort, with varying germline *A3B* deletion status: homozygous deleted (A3B^del/del^), homozygous wild type (A3B^WT/WT^), and heterozygous (A3B^del/WT^) samples. All the tumours were histologically classified as invasive ductal carcinoma (IDC) and had similar stages and grades. A total of 18 samples were screened. Histology quality checks were performed to ensure only samples with good morphology and cellularity were chosen for sequencing. All the selected samples had an average RIN (RNA Integrity Number) score of 8.24, indicating that the samples had highly intact RNA and would be suitable candidates for Stereo-seq profiling. There were 15 samples that passed the screening, and eight samples were chosen for the profiling, with three samples for A3B^del/del^, three samples for A3B^WT/WT^, and two samples for A3B^del/WT^. The A3Bdel status was previously determined in [11].

### Stereo-seq profling

The sectioned tissue slices were placed onto a Stereo-seq capture chip and fixed with methanol for 30 minutes at −20°C. The tissues were stained with Qubit ssDNA reagent (Invitrogen). After mounting with glycerol, the chips were imaged using a fluorescent microscope (Olympus). The tissues on the chips were processed according to manufacturer’s protocol (Stereo-seq Transcriptomics T Kit). In brief, the sections were first permeabilised using the Permeasbilisation Mix containing HCl (Sigma) for an optimised time that was determined using the Stereo-seq Permeabilisation Kit. Following reverse transcription, cDNA products were released from the chips using the cDNA Release Enzyme Mix. The cDNA mix was collected and purified by 0.8× beads (AMPure) selection, then amplified by the cDNA Amplification Mix with cDNA primers, and then further purified by 0.6× beads (AMPure). Purified cDNA samples were used for cDNA library preparation according to manufacturer’s protocol for Library Preparation Kit. The cDNA libraries were sequenced on a MGI DNBSEQ-T7 Sequencer (BGI).

### Stereo-seq data processing

Raw reads were processed using the SAW pipeline (https://github.com/STOmics/SAW). The final outputs were count matrices where each row represented a coordinate identity (CID), while columns were gene expression counts. Basic data processing was conducted using Stereopy (v. 1.4.0) in the Python environment (v. 3.8.19). The analysis was conducted at the BIN100 level as the sweet spot between bin quality and resolution. Ribosomal and mitochondrial genes were first filtered, then the bins were filtered for those with more than 20 transcripts and genes that were shared by at least 3 spots. The data was then normalised, log transformed and scaled in a similar process as Seurat’s SCTransform. Principal Component Analysis (PCA) was conducted with 30 principal components, followed by neighbourhood graph construction with 10 neighbours. Further dimensionality reduction was performed using Uniform Manifold Approximation Projection (UMAP). Lastly, clustering was performed with the Leiden algorithm for individual samples to retain their individual heterogeneity. The processed Stereopy object was exported as an AnnData object with the *st.io.write_h5ad function*, and was also converted into a Seurat object using the *st.io.stereo_to_anndata* function.

### Cell type deconvolution

Cell type deconvolution was carried out with RCTD [45] embedded within the *Spacexr* package (https://github.com/dmcable/spacexr) for each sample. A single-cell breast cancer atlas [46] was used as reference for the deconvolution using the “full” option. Only the cells originating from the TNBC samples were included in the Reference object. The Stereo-seq data was imported as a Seurat object and converted into a SpatialRNA object. RCTD generated an output matrix where each bin was given a weight for each of the cell types, which were treated as probability scores for subsequent analyses. The cell types weights were further aggregated into broader categories for more general analyses. The weights were grouped into major cell type categories and averaged across all bins. Hierarchical clustering of these samples was performed with Pearson correlation-based distance metric and average linkage using Numpy (v. 1.23.5), Scipy (v. 1.10.1), and the *clustermap* function from Seaborn (v. 0.12.2).

### Somatic variant calling

Somatic mutations were called from the mapped Stereo-seq reads using Mutect2, using previously conducted whole exome sequencing of buffy coat from the same patient as our matched normal sample. Only variants that passed FilterMutectCalls were then annotated with Funcotator. A series of filters that had been performed for RNA-seq-derived somatic mutations [47, 48] were applied to our data. The annotated variants went through a multi-step filtration process with four filters. Variants with high variant allele frequency (VAF) were found to likely be false positives, and the studies respectively removed variants with VAF more than 0.9 and 0.7, we settled on a median threshold of 0.8. We also excluded variants with TLOD score less than 5.6. We then removed variants identified as short tandem repeats (STR). Lastly, we excluded variants that were identified in blacklisted regions, which include sex chromosomes, mitochondrial chromosomes and unfinished scaffolds.

We validated the somatic variants using matched RNAseq and WES tumour data. For each sample, we filtered for somatic variants found in both RNAseq and WES tumour, and quantified the number of variants supporting each variant across Stereo-seq, RNAseq, and WES tumour (**Supp Table 1**). We manually examined the transcripts in IGV and identified variants with more than 10 transcripts in Stereo-seq and quantified variants with at least 1 allele supporting the mutation. Concordance was then calculated by taking the percentage of supported variants over the total number of overlapping variants (**Supp Table 2**). We further validated germline variants by randomly sampling 10 germline variants from each sample and examining them in IGV, comparing variant quality across WES normal, Stereo-seq, RNAseq, and WES tumour. For every germline variant identified in WES normal, transcripts covered in Stereo-seq and RNAseq also contained the mutations (**Supp Table 3**).

In order to visualise the spatial location of mutations, the coordinates of each variant had to be mapped back to its original transcript. After filtering the variants for SNVs of interest, transcripts with the corresponding chromosome and position were extracted from the mapped reads with Samjdk (https://lindenb.github.io/jvarkit/SamJdk.html), part of the *jvarkit* package written by Pierre Lindenbaum. The coordinates of each transcript were then extracted. Based on the resolution of BIN100, the coordinates were rounded accordingly to the nearest 100, and transcripts in the same bins were aggregated. The rounded SNV transcript counts were then normalised with respect to the total transcript counts for each bin to get a mutation score.

### Mutational signature analyses

Due to the low numbers of variants detected per bin, we chose to define and quantify mutational signatures in this study based on the most common trinucleotide context for each mutational signature as defined in the COSMIC library of mutational signatures. Scoring and visualization of each COSMIC mutational signature was based on the number of transcripts containing variants that had those trinucleotide contexts. Our mutational signature analyses should thus be considered a proxy for COSMIC mutational signature rather than a true quantification of the actual mutational signatures found in COSMIC.

### Neoantigens prediction

Using the annotated variants, additional filtering was performed to exclude variants in the HLA and immunoglobulin genes. Further annotations were carried out with Ensembl Variant Effect Predictor (VEP) (v. 112.0). Phasing, which resolves the genotypes uncovered by variant calling into haplotypes, was performed using WhatsHap [47] (v. 2.3). The phased variants were then finally annotated again with VEP. Neoantigens were predicted using pVACseq [49] (v. 4.1.0) with the VEP-annotated variants as the main input, tumour transcripts, as well as patient-specific HLA types that had been genotyped using HLA-HD [50] (v. 1.7.0). The phased option was fulfilled with the phased variant generated in the previous step. The algorithms used for the prediction included MHCflurry, MHCflurryEL, NNalign, NetMHC, NetMHCIIpan, NetMHCIIpanEL, NetMHCpan and NetMHCpanEL. The range for class I epitope length was between 8 to 11, while the length for class II epitope was 15. The predictions were patient-specific and a set of neoantigens were predicted for each sample. The same downstream processing steps as the somatic variants were repeated to visualise the spatial expression of the neoantigens.

### Statistical analyses

Statistical analyses were carried out using SciPy (v. 1.10.1) and annotated using statannotations (v. 0.7.2).

## Acknowledgements

Cancer Research Malaysia receives charitable funding from the Scientex Foundation, Estée Lauder Companies, Yayasan Petronas, and Yayasan Sime Darby, which contributed to the funding of this study.

## Ethics declaration

Patient recruitment and sample collection was reviewed and approved by the Independent Ethics Committee, Ramsay Sime Darby Health Care (Reference numbers: 201109.4, 201208.1, and 201805.2). Written informed consent to participation in research was given by each individual patient.

## References

1. Nik-Zainal, S. and S. Morganella, Mutational Signatures in Breast Cancer: The Problem at the DNA Level. Clin Cancer Res, 2017. 23(11): p. 2617–2629.

2. Dananberg, A., et al., APOBEC Mutagenesis in Cancer Development and Susceptibility. Cancers (Basel), 2024. 16(2).

3. Jamal-Hanjani, M., et al., Tracking the Evolution of Non-Small-Cell Lung Cancer. N Engl J Med, 2017. 376(22): p. 2109–2121.

4. Gupta, A., et al., APOBEC3 mutagenesis drives therapy resistance in breast cancer. Nat Genet, 2025. 57(6): p. 1452–1462.

5. Yang, Y., N. Liu, and L. Gong, An overview of the functions and mechanisms of APOBEC3A in tumorigenesis. Acta Pharm Sin B, 2024. 14(11): p. 4637–4648.

6. Butler, K. and A.R. Banday, APOBEC3-mediated mutagenesis in cancer: causes, clinical significance and therapeutic potential. J Hematol Oncol, 2023. 16(1): p. 31.

7. Kidd, J.M., et al., Population stratification of a common APOBEC gene deletion polymorphism. PLoS Genet, 2007. 3(4): p. e63.

8. Caval, V., et al., A prevalent cancer susceptibility APOBEC3A hybrid allele bearing APOBEC3B 3’UTR enhances chromosomal DNA damage. Nat Commun, 2014. 5: p. 5129.

9. Nik-Zainal, S., et al., Association of a germline copy number polymorphism of APOBEC3A and APOBEC3B with burden of putative APOBEC-dependent mutations in breast cancer. Nat Genet, 2014. 46(5): p. 487–91.

10. Pan, J.W., et al., The molecular landscape of Asian breast cancers reveals clinically relevant population-specific differences. Nat Commun, 2020. 11(1): p. 6433.

11. Pan, J.W., et al., Germline APOBEC3B deletion increases somatic hypermutation in Asian breast cancer that is associated with Her2 subtype, PIK3CA mutations and immune activation. Int J Cancer, 2021. 148(10): p. 2489–2501.

12. Lonsdale, J., et al., The Genotype-Tissue Expression (GTEx) project. Nature Genetics 2013 45:6, 2013. 45(6): p. 580–5.

13. Cescon, D.W., B. Haibe-Kains, and T.W. Mak, APOBEC3B expression in breast cancer reflects cellular proliferation, while a deletion polymorphism is associated with immune activation. Proc Natl Acad Sci U S A, 2015. 112(9): p. 2841–6.

14. Wen, W.X., et al., Germline APOBEC3B deletion is associated with breast cancer risk in an Asian multi-ethnic cohort and with immune cell presentation. Breast Cancer Research, 2016. 18(1): p. 56.

15. Yang, J., et al., Comprehensive Analyses Reveal Effects on Tumor Immune Infiltration and Immunotherapy Response of APOBEC Mutagenesis and Its Molecular Mechanisms in Esophageal Squamous Cell Carcinoma. Int J Biol Sci, 2023. 19(8): p. 2551–2571.

16. Faden, D.L., et al., APOBEC mutagenesis is tightly linked to the immune landscape and immunotherapy biomarkers in head and neck squamous cell carcinoma. Oral Oncol, 2019. 96: p. 140–147.

17. Huang, G., et al., APOBEC family reshapes the immune microenvironment and therapy sensitivity in clear cell renal cell carcinoma. Clin Exp Med, 2024. 24(1): p. 212.

18. Sarangi, P., Role of indoleamine 2, 3-dioxygenase 1 in immunosuppression of breast cancer. Cancer Pathog Ther, 2024. 2(4): p. 246–255.

19. Melssen, M.M., et al., Barriers to immune cell infiltration in tumors. J Immunother Cancer, 2023. 11(4).

20. Chung, S.W., Y. Xie, and J.S. Suk, Overcoming physical stromal barriers to cancer immunotherapy. Drug Deliv Transl Res, 2021. 11(6): p. 2430–2447.

21. Cervantes-Gracia, K., et al., APOBECs orchestrate genomic and epigenomic editing across health and disease. Trends Genet, 2021. 37(11): p. 1028–1043.

22. Niocel, M., et al., The DNA damage induced by the Cytosine Deaminase APOBEC3A Leads to the production of ROS. Sci Rep, 2019. 9(1): p. 4714.

23. Xia, L., X. Yan, and H. Zhang, Mitochondrial DNA-activated cGAS-STING pathway in cancer: Mechanisms and therapeutic implications. Biochim Biophys Acta Rev Cancer, 2025. 1880(1): p. 189249.

24. Kim, J., H.S. Kim, and J.H. Chung, Molecular mechanisms of mitochondrial DNA release and activation of the cGAS-STING pathway. Exp Mol Med, 2023. 55(3): p. 510–519.

25. Li, T. and Z.J. Chen, The cGAS-cGAMP-STING pathway connects DNA damage to inflammation, senescence, and cancer. J Exp Med, 2018. 215(5): p. 1287–1299.

26. Hu, A., et al., Harnessing innate immune pathways for therapeutic advancement in cancer. Signal Transduct Target Ther, 2024. 9(1): p. 68.

27. Liu, T., et al., NF-kappaB signaling in inflammation. Signal Transduct Target Ther, 2017. 2(1): p. 17023-.

28. Abdou, Y., et al., Immunotherapy in triple negative breast cancer: beyond checkpoint inhibitors. NPJ Breast Cancer, 2022. 8(1): p. 121.

29. Denkert, C., et al., Tumour-infiltrating lymphocytes and prognosis in different subtypes of breast cancer: a pooled analysis of 3771 patients treated with neoadjuvant therapy. Lancet Oncol, 2018. 19(1): p. 40–50.

30. He, T.F., et al., Multi-panel immunofluorescence analysis of tumor infiltrating lymphocytes in triple negative breast cancer: Evolution of tumor immune profiles and patient prognosis. PLoS One, 2020. 15(3): p. e0229955.

31. Kim, G., et al., The Contribution of Race to Breast Tumor Microenvironment Composition and Disease Progression. Front Oncol, 2020. 10: p. 1022.

32. Ding, S., et al., Single-cell atlas reveals a distinct immune profile fostered by T cell-B cell crosstalk in triple negative breast cancer. Cancer Commun (Lond), 2023. 43(6): p. 661–684.

33. Wang, X., et al., Spatial transcriptomics reveals substantial heterogeneity in triple-negative breast cancer with potential clinical implications. Nat Commun, 2024. 15(1): p. 10232.

34. Smith, N.J., et al., Differentiation signals induce APOBEC3A expression via GRHL3 in squamous epithelia and squamous cell carcinoma. EMBO J, 2025. 44(1): p. 1–29.

35. Burns, M.B., et al., APOBEC3B is an enzymatic source of mutation in breast cancer. Nature, 2013. 494(7437): p. 366–70.

36. Chan, K., et al., An APOBEC3A hypermutation signature is distinguishable from the signature of background mutagenesis by APOBEC3B in human cancers. Nat Genet, 2015. 47(9): p. 1067–72.

37. Alexandrov, L.B., et al., The repertoire of mutational signatures in human cancer. Nature, 2020. 578(7793): p. 94–101.

38. Alexandrov, L.B., et al., Signatures of mutational processes in human cancer. Nature, 2013. 500(7463): p. 415–21.

39. Nik-Zainal, S., et al., Landscape of somatic mutations in 560 breast cancer whole-genome sequences. Nature, 2016. 534(7605): p. 47–54.

40. Zhao, L., et al., Tertiary lymphoid structures in diseases: immune mechanisms and therapeutic advances. Signal Transduct Target Ther, 2024. 9(1): p. 225.

41. Narvaez, D., et al., The Emerging Role of Tertiary Lymphoid Structures in Breast Cancer: A Narrative Review. Cancers (Basel), 2024. 16(2).

42. Wang, Q., et al., Heterogeneity of tertiary lymphoid structures predicts the response to neoadjuvant therapy and immune microenvironment characteristics in triple-negative breast cancer. Br J Cancer, 2025. 132(3): p. 295–310.

43. Boissiere-Michot, F., et al., Prognostic value of tertiary lymphoid structures in triple-negative breast cancer: integrated analysis with the tumor microenvironment and clinicopathological features. Front Immunol, 2024. 15: p. 1507371.

44. Tan, M.-M., et al., A case-control study of breast cancer risk factors in 7,663 women in Malaysia. PLoS ONE, 2018. 13(9): p. e0203469.

45. Cable, D.M., et al., Robust decomposition of cell type mixtures in spatial transcriptomics. Nature Biotechnology, 2022. 40(4): p. 517–526.

46. Wu, S.Z., et al., A single-cell and spatially resolved atlas of human breast cancers. Nat Genet, 2021. 53(9): p. 1334–1347.

47. Martin, M., P. Ebert, and T. Marschall, *Read-Based Phasing and Analysis of Phased Variants with WhatsHap*, in Methods in Molecular Biology. 2023, Springer US. p. 127–138.

48. Garcia-Nieto, P.E., A.J. Morrison, and H.B. Fraser, The somatic mutation landscape of the human body. Genome Biol, 2019. 20(1): p. 298.

49. Hundal, J., et al., pVAC-Seq: A genome-guided in silico approach to identifying tumor neoantigens. Genome Medicine, 2016. 8(1).

50. Kawaguchi, S., et al., HLA-HD: An accurate HLA typing algorithm for next-generation sequencing data. Hum Mutat, 2017. 38(7): p. 788–797.

