## Supplementary material for "Spatial patterns of APOBEC mutagenesis in the tumour microenvironment of Asian breast cancer": Supp Figures 1-10

Tan, ZC et al.

**Supplementary information**

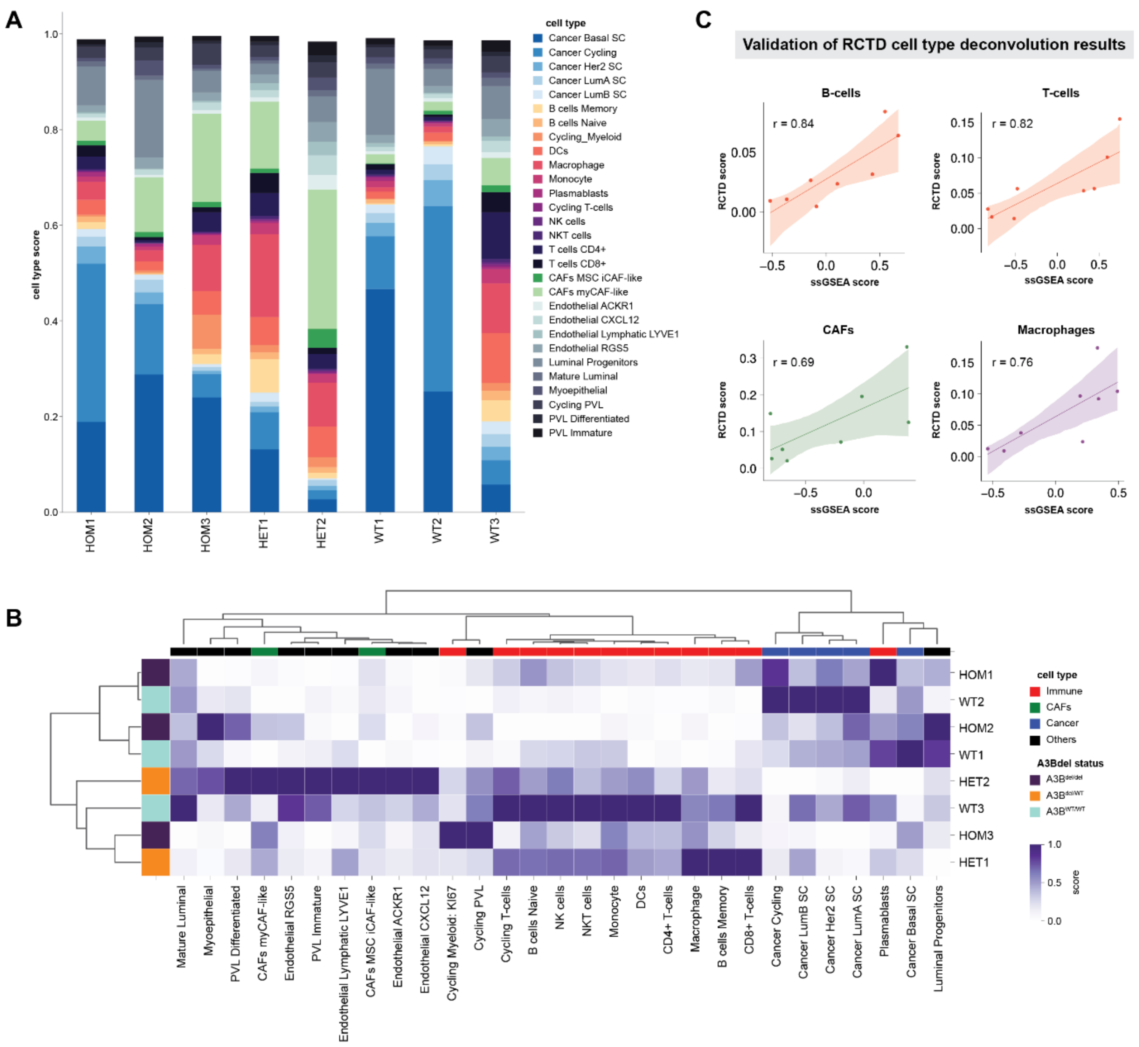


**Supp Figure 1**. Expanded minor cell type analysis and validation of deconvolution results. A: Composition of tumour microenvironments of each TNBC sample subsetted to the minor cell type resolution. B: Unsupervised hierarchical clustering of minor cell types across samples. The scores for each sample were first adjusted to a total of 1 prior to clustering. C: Pearson correlation between ssGSEA scores from matched bulk RNA-seq samples and pseudobulk cell type scores from the Stereo-seq samples, as validation for our cell type deconvolution results.

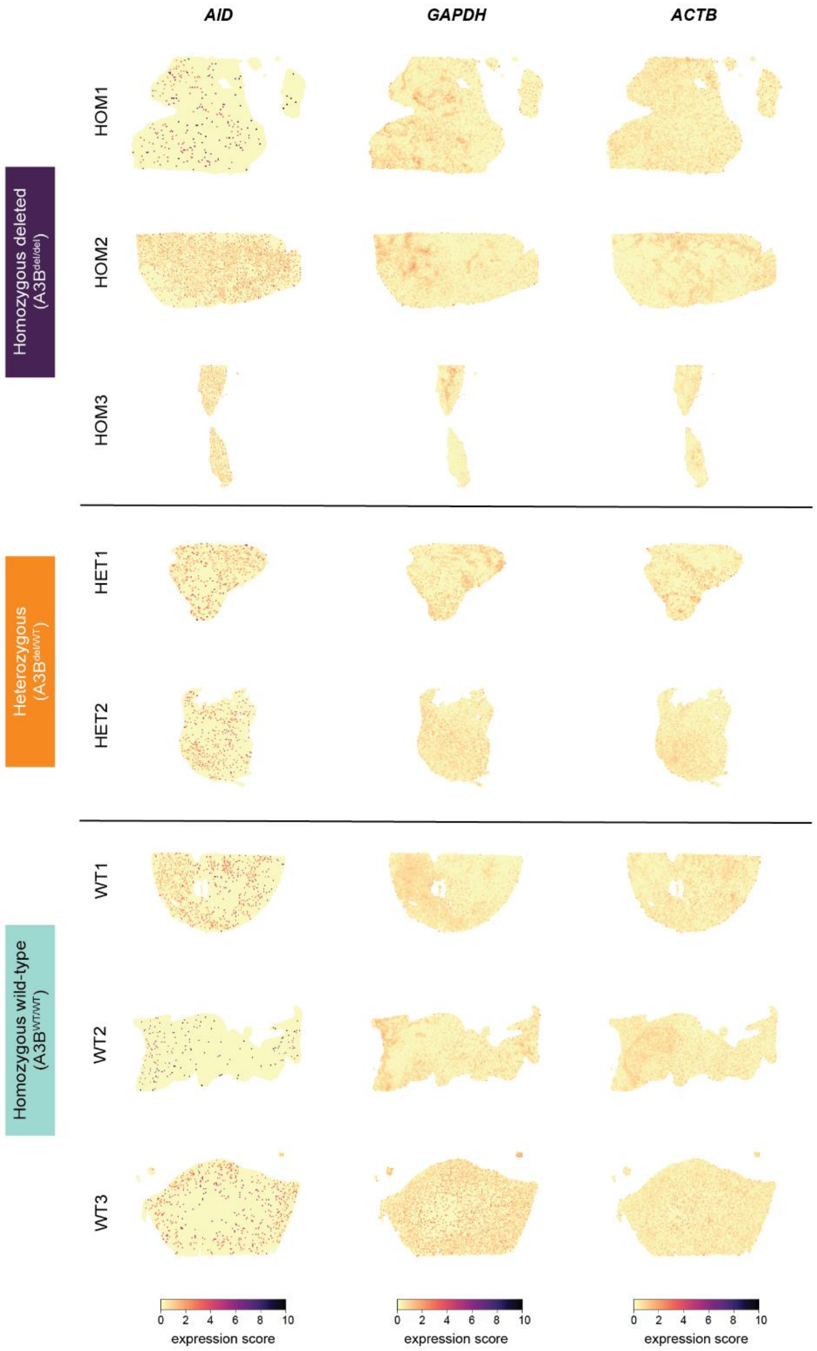


**Supp Figure 2**. Spatial expression patterns of housekeeping genes: *AID*, *GAPDH*, *ACTB*.

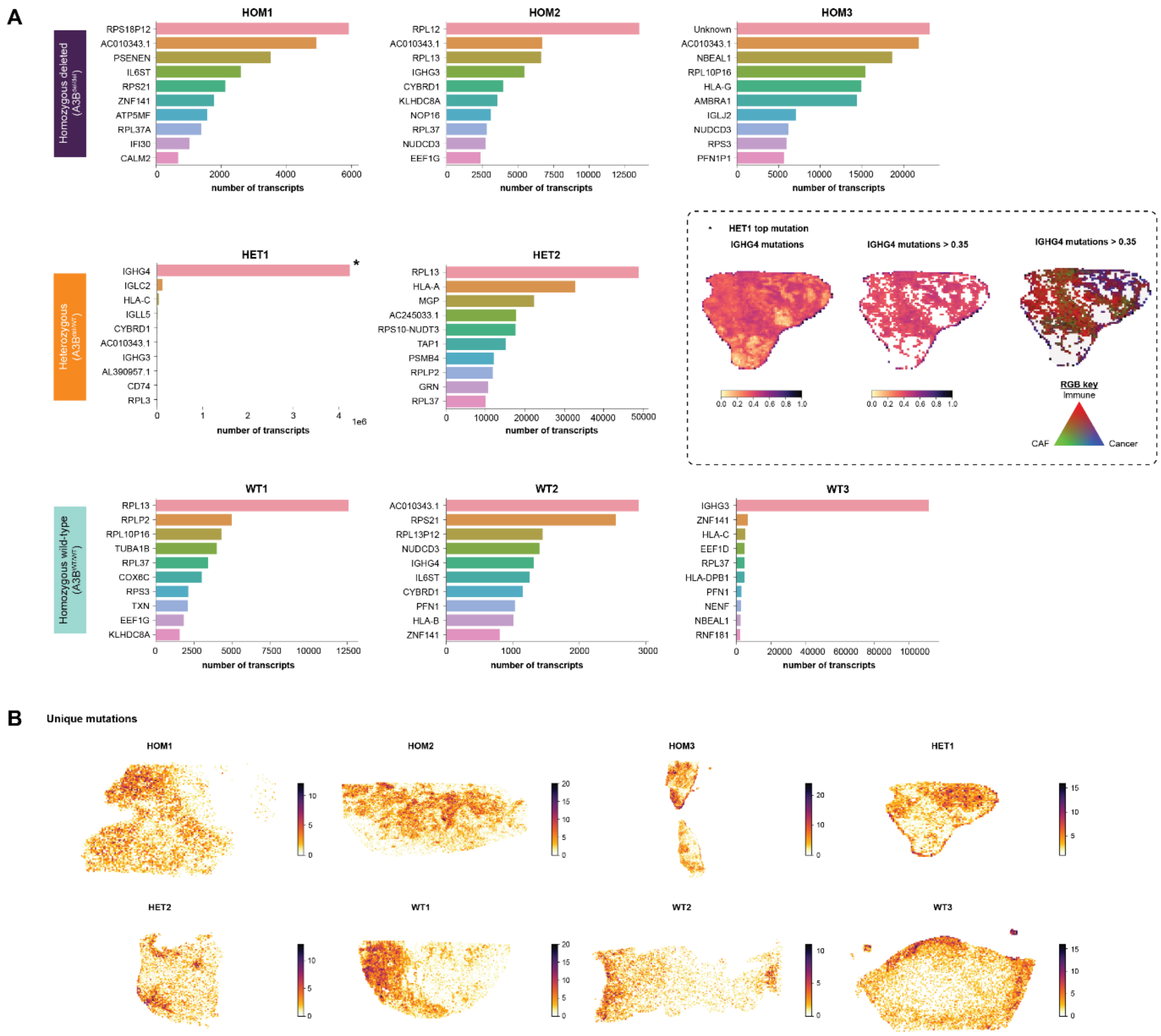


**Supp Figure 3**. Somatic single nucleotide variants (SNVs) discovered from the TNBC cohort. A: Quantification of somatic gene mutations across samples, prior to mapping to individual bins and normalisation. Barplots display the top 10 mutations in each sample. Dotted box shows the spatial expression of *IGHG4*, the top mutation in HET1, followed by regions with high mutation scores, and the cellular composition context of those regions. B: Spatial tumour mutational burden (spatial TMB) by quantifying the number of unique SNVs in each bin.

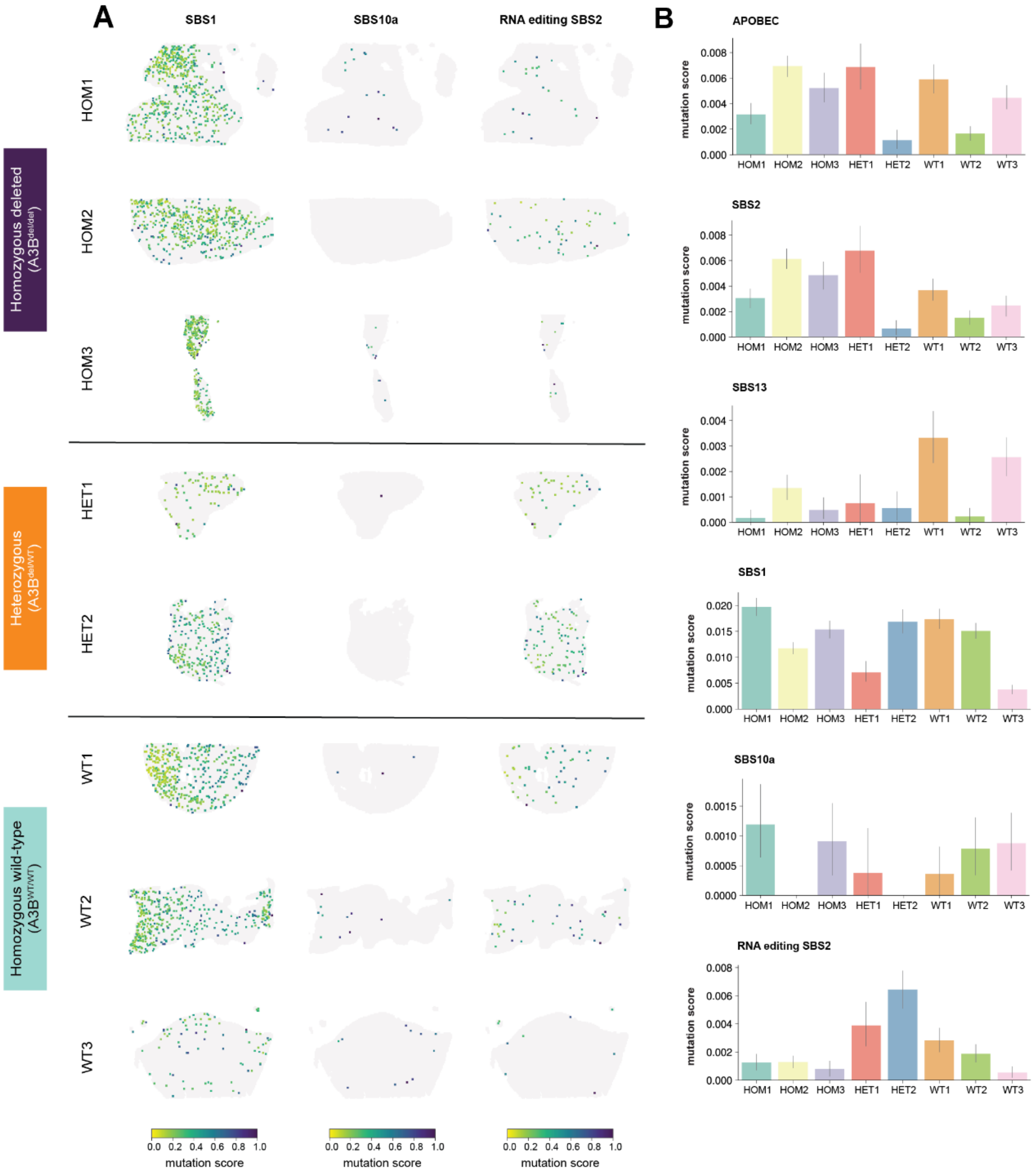


**Supp Figure 4**. Distribution of mutational signatures. A: Spatial distribution of COSMIC mutational signatures SBS1, SBS10a, and RNA-editing SBS2. B: Average mutation scores for APOBEC, SBS2, SBS13, SBS1, SBS10a, and RNA-editing SBS2 mutations.

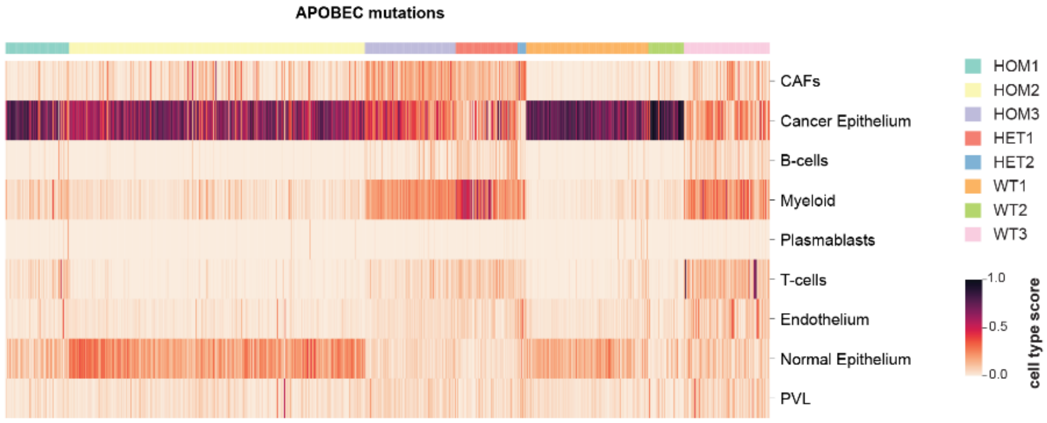


**Supp Figure 5**. Heatmap showing major cell type scores for bins predicted of having APOBEC mutations across all samples.

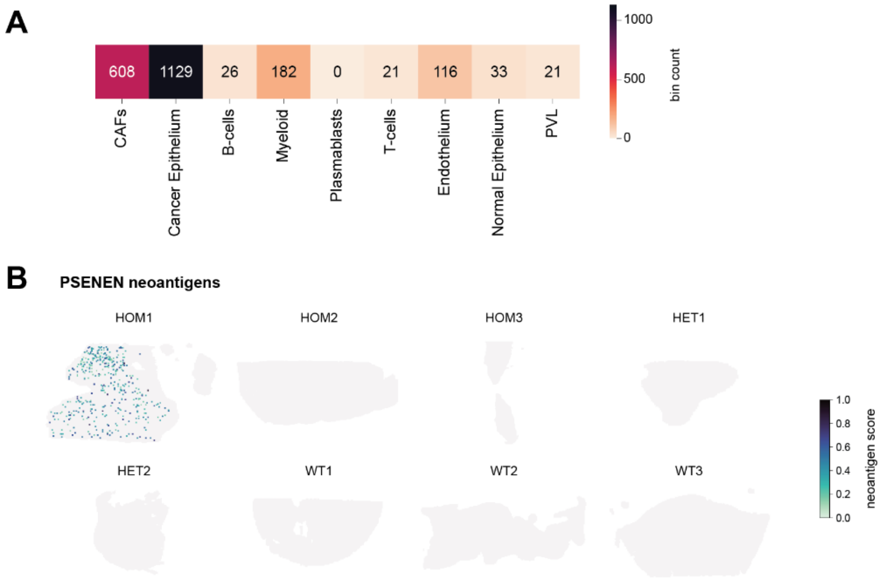


**Supp Figure 6**. Quantification and distribution of candidate neoantigens. A: Quantification of bins predicted to contain neoantigens. Each bin was assigned a cell type based on the highest score prior to analysis. B: Spatial distribution of PSENEN neoantigen candidates across samples.

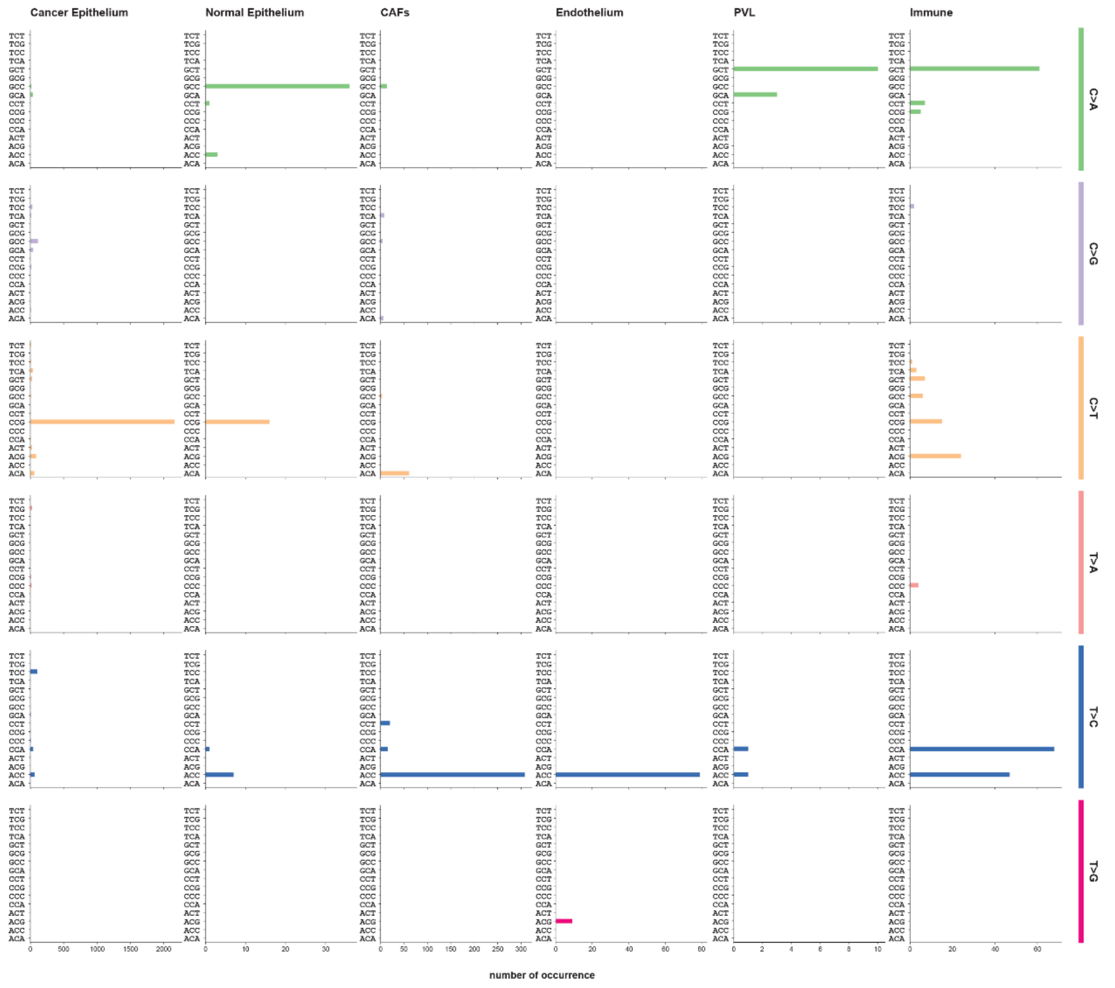


**Supp Figure 7**. Quantification of single base substitutions that are predicted give rise to neoantigens, limited to bins previously determined to be cancer epithelium, normal epithelium, CAFs, endothelium, PVL, and immune cells. The scale is not truncated and not share between plots.

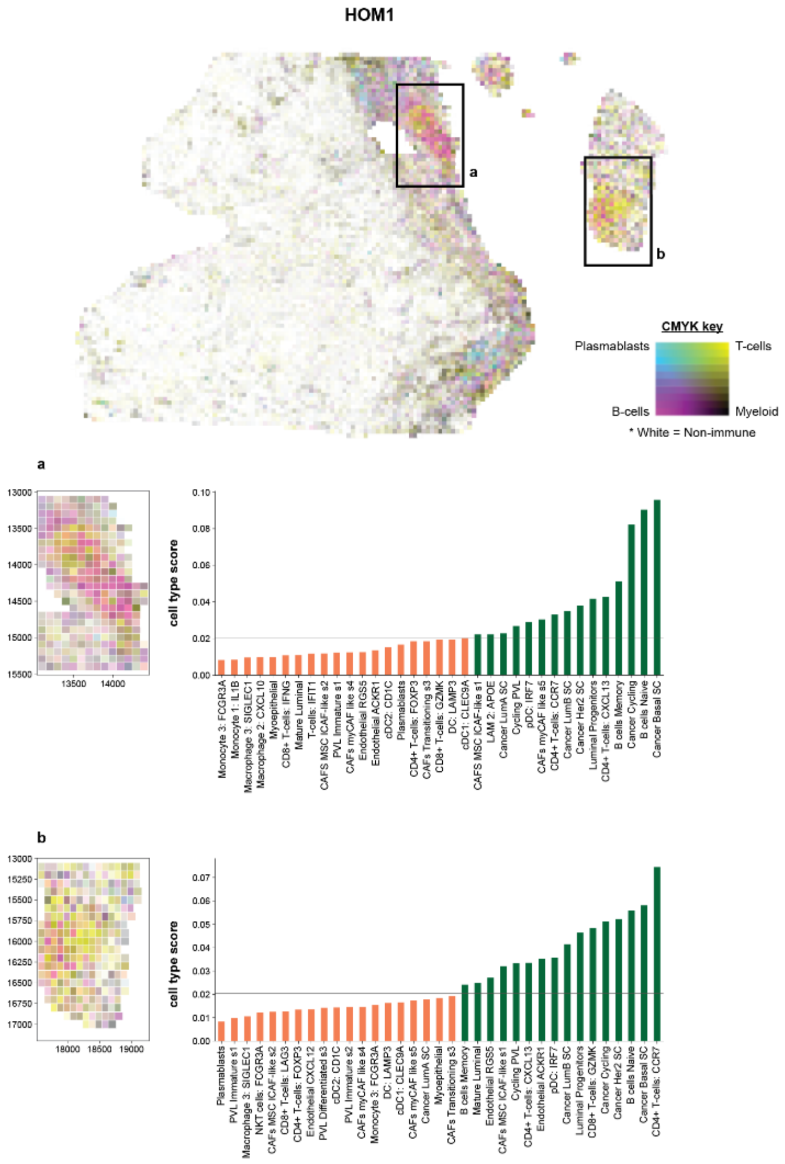


**Supp Figure 8**. Closer look at regions (a, b) suspected to be tertiary lymphoid structures (TLS) within HOM1. Detailed cellular composition for each region is shown, cell types exceeding the mean (horizontal line) are highlighted in green.

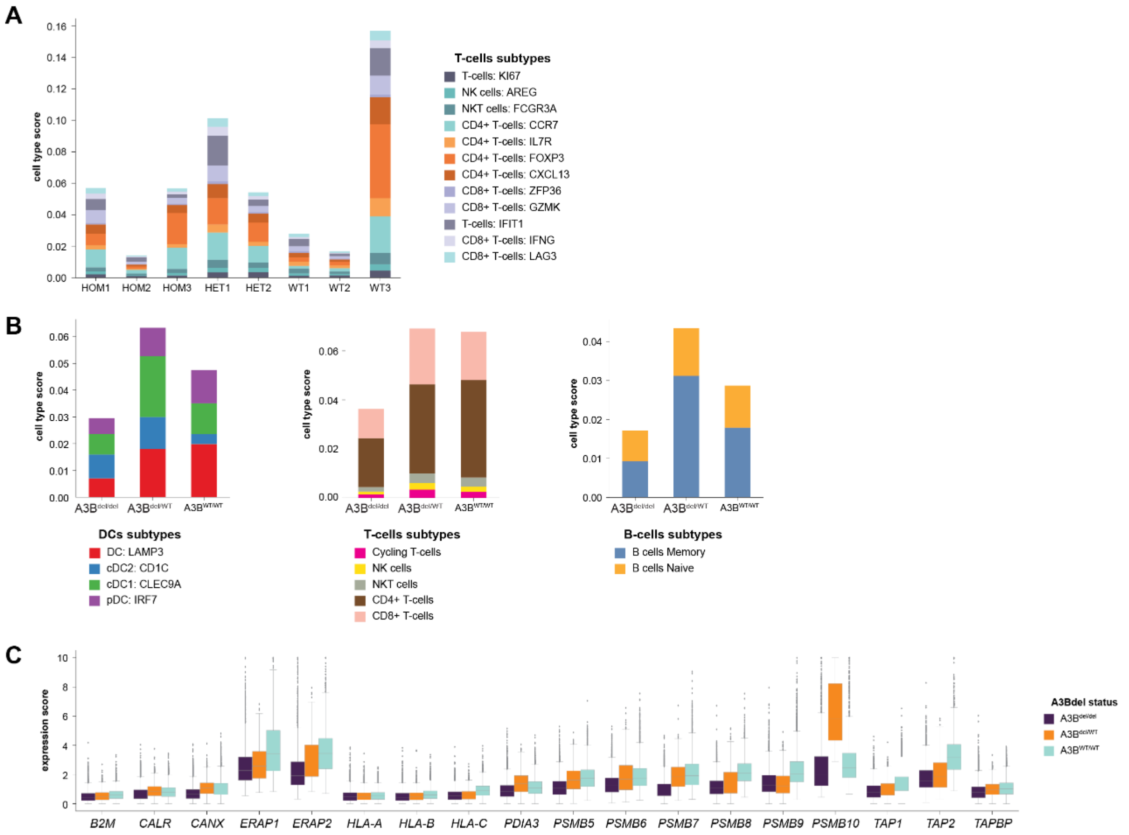


**Supp Figure 9**. Distribution of immune subtypes. A: Barplot showing the average cell type scores for T-cell subtypes across all samples. B: Barplots showing the average cell type scores for DCs, T-cells, and B-cells, each broken down into individual subtypes, across different germline *A3B* deletion statuses. C: Boxplots showing the distribution of non-zero gene expression scores for genes in the antigen presentation machinery across different germline *A3B* deletion statuses.

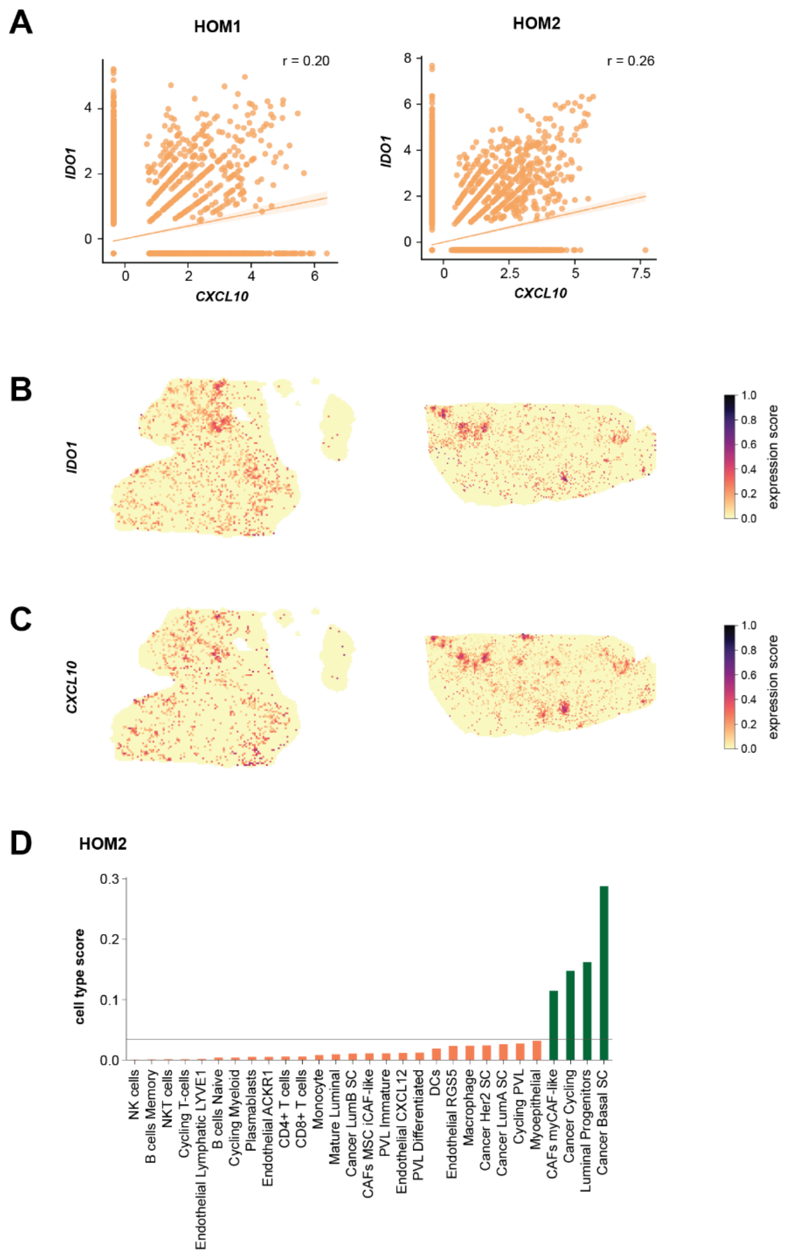


**Supp Figure 10**. Relationship between *IDO1* and *CXCL10* in HOM1 and HOM2. A: Pearson correlation between *IDO1* and *CXCL10* in HOM1 and HOM2, respectively. B: Spatial expression of *IDO1* in HOM1 and HOM2. C: Spatial expression of *CXCL10* in HOM1 and HOM2. D: Detailed cellular composition for HOM2, cell types exceeding the mean (horizontal line) are highlighted in green.
